# Dynamic Fading Memory and Expectancy Effects in Monkey Primary Visual Cortex

**DOI:** 10.1101/2023.11.06.565858

**Authors:** Yang Yiling, Johanna Klon-Lipok, Katharine Shapcott, Andreea Lazar, Wolf Singer

**Author notes:** Corresponding author: Wolf Singer, Prof. Dr. Ernst Strüngmann Institut gGmbH Deutschordenstraße 46, 60528 Frankfurt am Main Germany, Internet: https://www.esi-frankfurt.de/people/wolfsinger.

## Abstract

In order to investigate the involvement of primary visual cortex (V1) in working memory (WM), parallel, multisite recordings of multiunit activity were obtained from monkey V1 while the animals performed a delayed match-to-sample (DMS) task. During the delay period, V1 population firing rate vectors maintained a lingering trace of the sample stimulus that could be reactivated by intervening impulse stimuli that enhanced neuronal firing. This fading trace of the sample did not require active engagement of the monkeys in the DMS task and likely reflects the intrinsic dynamics of recurrent cortical networks in lower visual areas. This renders an active, attention-dependent involvement of V1 in the maintenance of working memory contents unlikely. By contrast, population responses to the test stimulus depended on the probabilistic contingencies between sample and test stimuli. Responses to tests that matched expectations were reduced which agrees with concepts of predictive coding.

## Introduction

Working memory (WM) refers to the ability to maintain and manipulate information in the absence of input. WM has traditionally been attributed to higher-order cortical areas, in particular prefrontal cortex (Funahashi et al., 1989; Fuster and Alexander, 1971; Kubota and Niki, 1971) and more recently to cooperative processes across multiple brain areas (Brincat et al., 2021; de Vries et al., 2020; Reinhart and Nguyen, 2019). There is also evidence for a recruitment of primary sensory areas like V1 in visual WM processes (D’Esposito, 2007; D’Esposito and Postle, 2015; Lara and Wallis, 2015; Pasternak and Greenlee, 2005; Scimeca et al., 2018; Serences, 2016; Sreenivasan et al., 2014; Supèr et al., 2001; van Kerkoerle et al., 2017). For example, information held in visual WM can be decoded from V1 activity (Christophel et al., 2018; Emrich et al., 2013; Ester et al., 2013; Ester et al., 2009; Harrison and Tong, 2009; Lawrence et al., 2018; Lorenc et al., 2018; Rademaker et al., 2019; Serences et al., 2009; Wolff et al., 2015; Wolff et al., 2017); the volume (Bergmann et al., 2016) and activity (Ester et al., 2013; Iamshchinina et al., 2021a, b) of V1 are positively correlated with behavioural performance in WM tasks. Early evidence suggests that WM is mediated by persistent firing during the delay period (Constantinidis et al., 2018; Funahashi et al., 1989; Fuster and Alexander, 1971; Haller et al., 2018; Kaminski et al., 2017; Kornblith et al., 2017; Kubota and Niki, 1971). However, this view has been contested (Lundqvist et al., 2018a) because WM contents were decodable only from short, temporally segregated bouts of activity (Lundqvist et al., 2018b; Lundqvist et al., 2016; Romo et al., 1999). Other evidence suggests the existence of covert, activity-independent traces (Erickson et al., 2010; Fiebig and Lansner, 2017; Mongillo et al., 2008; Rose et al., 2016; Stokes, 2015; Sugase-Miyamoto et al., 2008; Trubutschek et al., 2019; Wolff et al., 2017) that can be activated by “pinging” the brain with unspecific stimuli (Wolff et al., 2015; Wolff et al., 2017) or transcranial magnetic stimulation (Rose et al., 2016).

However, most of these studies used neuroimaging techniques in humans in order to retrieve the traces of WM contents, which limits the identification of the underlying neuronal signals. Thus, it is unclear whether the signals recorded from V1 that permit decoding of WM contents reflect reverberating activity (“fading memory”) within the recurrent networks of lower visual areas (Buonomano and Maass, 2009; Nikolić et al., 2009), or result from top down projections that involve V1 in WM. Therefore, we set out to investigate at the neuronal level whether WM contents can be decoded from neuronal population activity in V1, whether pinging could revive WM traces, and whether persistent information about the stimulus could be attributed to fading memory in local circuits or showed the attention- and task-dependent properties of WM related activity.

Another goal of the present study was to investigate whether V1 responses are shaped by priors stored in memory. Responses of V1 neurons to external stimuli depend on both stimulus features and internal priors (Aitchison and Lengyel, 2017; de Lange et al., 2018; Rao and Ballard, 1999; Singer, 2021). Stimuli matching priors of Gestalt principles modify the synchronization patterns (Gray et al., 1989; Gray and Singer, 1989), correlation structure (Bányai et al., 2019), sequential activation (Yiling et al., 2023) and response amplitude (Kapadia et al., 1995) of neuronal responses. Spatial predictability derived from stimulus context reduces firing rate and/or enhances oscillatory synchronization among neurons in V1 (Gray et al., 1989; Gray and Singer, 1989; Peter et al., 2019; Uran et al., 2022; Vinck and Bosman, 2016; Vinje and Gallant, 2000) and temporal predictability of stimuli suppresses V1 activation in humans (Alink et al., 2010; Kok et al., 2012). These observations support the notion that the visual system learns the statistical regularities in sensory input to optimize its responses (de Lange et al., 2018; Singer, 2021; Singer and Lazar, 2016).

To pursue the above goals we performed parallel multisite electrophysiological recording in awake monkey V1 while the animal performed a delayed match-to-sample (DMS) task. To test the possibility that WM content could serve as prior, we introduced probabilistic associations between stimulus-pairs in the DMS task in order to establish implicit predictions and investigated whether such priors modified V1 responses. We found that V1 population activity maintained an intrinsic, latent trace of the sample stimuli regardless of the need to actively engage WM. This trace could be reactivated and strengthened by an irrelevant, non-specific stimulus (Wolff et al., 2015; Wolff et al., 2017). Once probabilistic priors were established for sample-test pairs, V1 responses to predicted test stimuli were reduced.

## Results

We trained two monkeys to perform a DMS task that required stimulus encoding, retention of stimulus identity in working memory (WM) and a manual forced choice response. During the task, the monkey fixated a spot at the centre of the screen. Two stimulus images (“sample” and “test”) were presented sequentially for 500 ms each, separated by a delay period of 1500 ms (Figure 1a). The animal had to report whether the two stimuli were the same (“match”) or different (“nonmatch”), by pushing a mechanical lever forward or backward, respectively. The numbers of match and nonmatch trials were balanced in order not to bias the animal’s behavioural response. Stimuli were standardized images (Brodeur et al., 2010; Brodeur et al., 2014) of simple objects displayed on a grey screen (Figure 1b). A set of three images were used in each session (counterbalanced for sample and test positions), and the set of images varied between sessions. The position and size of the stimuli were tailored for each monkey to cover the ensemble of the receptive fields (RF) of the respective recording sites. For Monkey 1 (Figure 1b), stimuli subtended 4.4° of visual angle, and their centre was 2.36° lateral to the vertical meridian and 1.34° below the horizontal meridian. For Monkey 2, stimuli subtended 7.84° of visual angle, and their centre was 4.05° lateral to the vertical meridian and 2.70° below the horizontal meridian. On average, Monkey 1 performed 78.1 ± 1.7 % (s.e.m., n = 6 sessions) correct responses (Supplementary figure 1). The average reaction time for correct responses was 632.0 ± 174.7 ms (s.d., median 586.4 ms. Supplementary figure 1). The RF positions and the behavioural performance for Monkey 2 are shown in Supplementary figure 2. Throughout the paper analyses were always performed separately for each animal.

**Figure 1.**
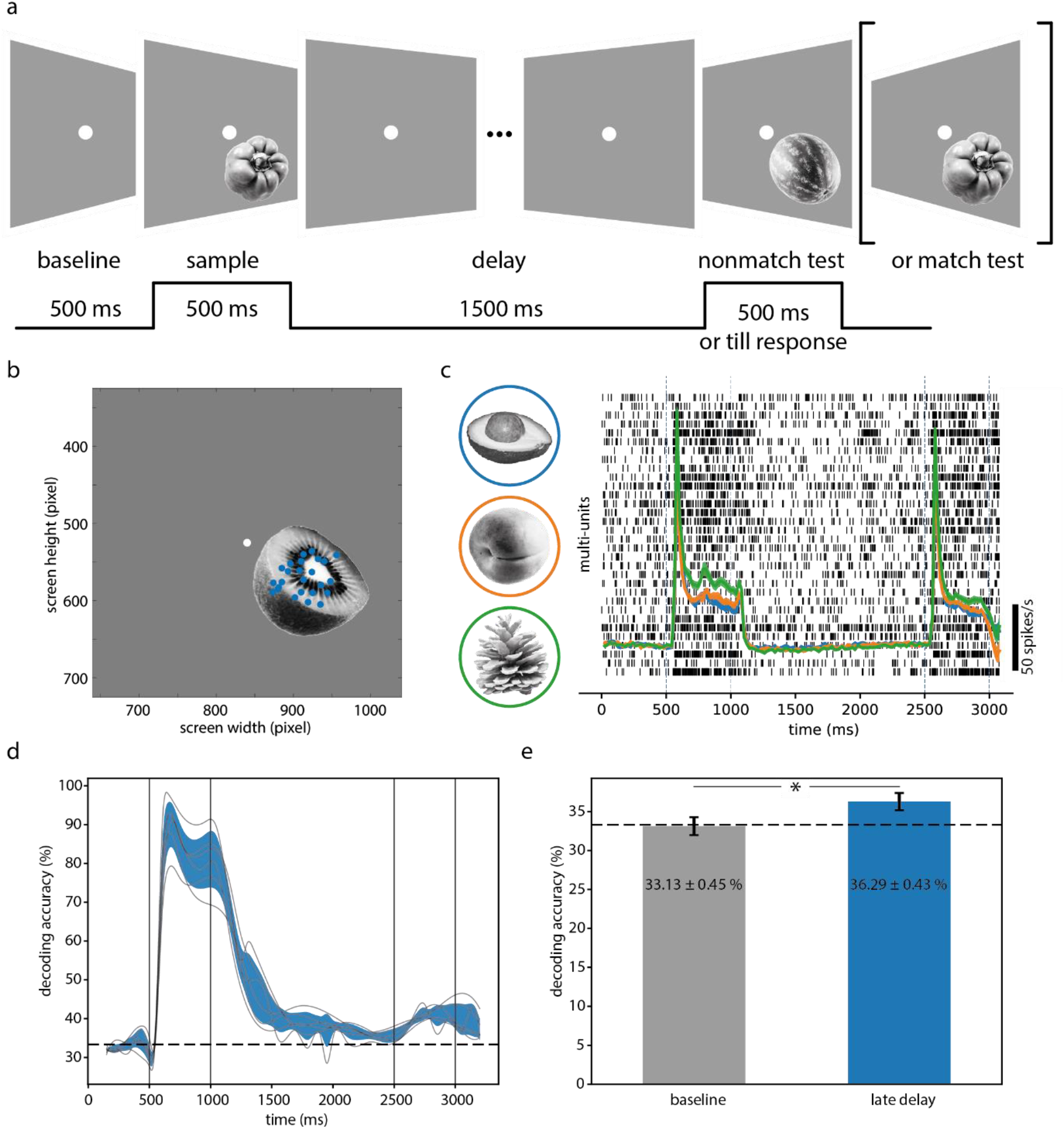
Robust trace of stimulus-specific information during the delay period. (a) Task structure and trial time course. (b) Positions of the stimulus, fixation point (white dot) and receptive fields (blue dots). (c) Raster plot of multi-unit activity in a single trial, overlaid with average population multi-unit firing rates for the three demo stimuli in match trials. Shades denote 95% confidence interval (barely visible due to large number of trials). (d) Time-resolved accuracy of decoding sample stimulus identity based on firing rate vectors. Grey traces: results from individual sessions; blue shades: 95% confidence interval around session average (n=6). (e) Comparison of decoding accuracies between late delay period (2000 – 2500ms) and pre-stimulus baseline (0 – 500ms). Error bars: 95% confidence intervals. Error numbers in legends denote standard error of the mean.

### Fading trace of sample stimulus

Multiunit activity (MUA) was recorded from visual area V1 (left hemisphere) with a chronically implanted 32-channel microdrive system (Gray Matter Research, Bozeman, Montana, USA) while the animal performed the DMS task. The increased firing of neurons evoked by the sample stimulus rapidly decayed during the WM delay period to the pre-stimulus baseline level or even below (Figure 1c). To compare the activity levels between the delay and baseline periods, we measured the spike counts (window size = 300 ms) for each channel and stimulus (pooled across sessions) at four time intervals in the delay period, and compared the spike counts with those in the baseline period preceding the sample stimulus (Supplementary figure 3 for Monkey 1, Supplementary figure 4 for Monkey 2). Spiking activity dropped below baseline level after the offset of the sample stimulus (1300 – 1600 ms: *t* = 13.91, *p* < 0.01; 1600 – 1900 ms: *t* = 15.64, *p* < 0.01; 1900 – 2200 ms: *t* = 7.51, *p* < 0.01. Paired *t*-test, Monkey 1. Statistics for Monkey 2 in Supplementary figure 4), but recovered to a level slightly above baseline towards the end of the delay period (2200 – 2500 ms: *t* = –3.24, *p* = 0.00014). There was no clear indication for a sustained elevation of discharge rates during the delay period.

To examine the amount of stimulus-specific information in the population vector of responses to the sample stimulus, we trained decoders (linear discriminant analysis, LDA) at successive time points to predict the sample stimulus identity from the population firing rate vector (window size 100 ms, step size 50 ms). The decoding accuracy (Figure 1d and Supplementary figure 4b) was highest during stimulus presentation (500 – 1000 ms), decayed after stimulus offset but remained above chance level (33.3%, 3 stimuli per session) throughout the delay period. The average decoding accuracy (36.29 ± 0.43 %, s.e.m., n = 6 sessions) for the sample stimulus in the last 500 ms of the delay period (from 2000 to 2500 ms) was still significantly above chance level (*t* = 6.26, *p* = 0.00153, *t*-test, two-sided unless noted otherwise), and higher than the baseline level (33.13 ± 0.45 %, s.e.m., *t* = –3.68, *p* = 0.0142, paired *t*-test) which did not differ from chance (*t* = –0.42, *p* = 0.691). Similar results were obtained from three other sets of experiments which used different numbers of stimuli (Supplementary figure 5a), and also from Monkey 2 (Supplementary figure 4b-c). To test further the robustness of the findings against the variation of stimuli, in a separate set of experiments we used gratings as stimuli in the same DMS task. Here, the animal was required to discriminate the orientations of the sample vs. test gratings (spatial frequency 3 cycles per degree; 4 non-cardinal orientations per session; 6 sessions in total). Interestingly, in this set of experiments, the trailing sample stimulus information decayed rapidly to the baseline level (Supplementary figure 5b). However, in this DMS task with grating stimuli, the animal’s performance (65.6 ± 1.5 %, n = 6) was worse (*t* = 5.54, *p* < 0.01) than in the standard DMS task (78.1 ± 1.7%, n = 6), although still above chance level (*t* = 10.48, *p* < 0.001). As the simple grating stimuli could give rise to retinal afterimages, these results suggest that the trailing traces of the natural sample stimuli were not due to retinal adaptation. Thus, despite the low firing rates that did not differ much from baseline towards the end of the delay period, the population activity retained a low but robust trace of sample stimulus-specific information in the experiments where natural objects were used as stimuli.

To test whether the trailing stimulus-specific information was actually related to the WM task, we performed control experiments with passive viewing on an animal naive to the DMS task. Here, the two stimuli were shown at the same time points as in the DMS task, but the monkey was only required to attend to the fixation spot and was rewarded for detecting and responding to a colour change of the fixation spot (Methods). The monkey had to push the lever forward or backward, if the fixation point colour changed to green or blue, respectively. Nevertheless, stimulus-specific information about the irrelevant “sample” stimulus persisted throughout the delay interval (Supplementary figure 6), suggesting that the trailing stimulus information was not caused by the requirement to memorize the sample stimulus.

### Reactivation of latent memory by impulse stimulus

Modelling (Mongillo et al., 2008) and neuroimaging (Wolff et al., 2017) studies reported that a global, unspecific activation of neuronal networks can reveal latent or covert traces of information held in memory. To examine whether such a manipulation could enhance sample-specific information during the delay interval, we modified the DMS task and inserted a full-screen impulse stimulus (100 ms duration, 100% intensity) in the delay interval (Figure 2a). To indicate to the monkey that this intervening stimulus was irrelevant for the task we presented different types of impulse stimuli: linear gratings (2 sessions with 0° orientation, 2 sessions with 90° orientation), a concentric grating (1 session), or a blank white stimulus (1 session). We also applied these stimuli in the passive viewing tasks, presenting one of the four stimuli in four different sessions. These stimuli were applied at either 500 ms or 1000 ms after the offset of the sample stimulus (i.e., 1500 ms or 2000 ms in a trial. Figure 2b). We then performed the same time-resolved decoding analysis as described above, pooling the results across all the different sessions (n = 10). As shown in Figure 2b (Supplementary figure 7 for Monkey 2), decoding accuracy increased about 100 ms after these impulse stimuli and remained enhanced for 100 – 200 ms (at 1700 ms: accuracy with impulse 43.11 ± 1.02 % s.e.m., accuracy without impulse 35.38 ± 1.01 %, *t* = 4.16, *p* = 0.0024; at 2200 ms: accuracy with impulse 39.63 ± 0.52 %, accuracy without impulse 34.25 ± 0.82 %, *t* = 4.79, *p* = 0.00099, n = 10 sessions). The increase in decodability resembled closely in time the transient firing rate increase evoked by the impulse (Figure 2b).

**Figure 2.**
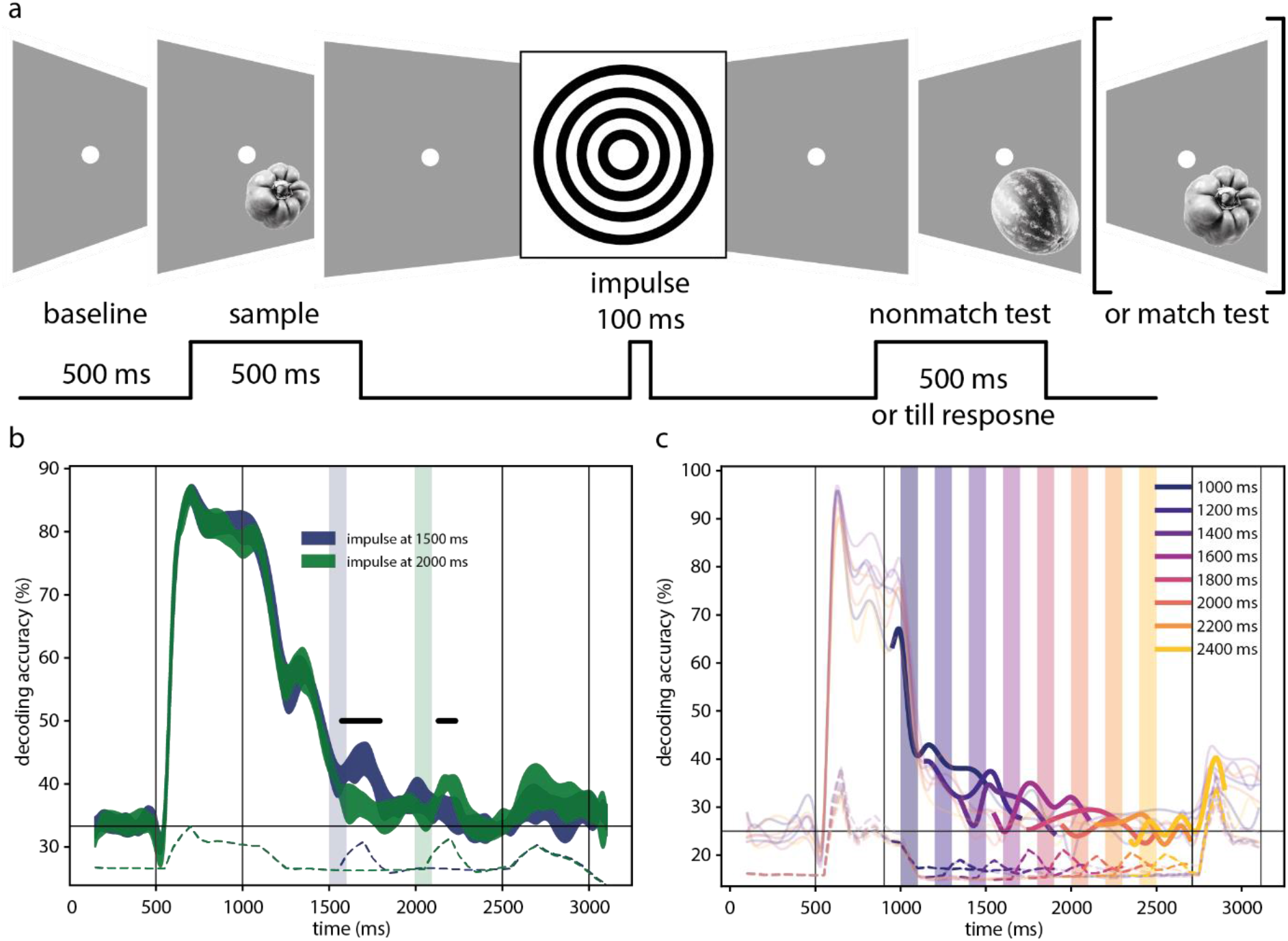
Visual impulse stimulus enhances latent memory trace. (a) Modified DMS trial structure. A full screen impulse stimulus (100 ms) is inserted in the delay period. (b) The accuracy of decoding the sample stimuli for two impulse conditions (blue: impulse at 1500 ms; green: impulse at 2000 ms). Shaded areas indicate 95% confidence intervals. Vertical blue and green bars mark the positions of the impulse stimuli at 1500 ms and 2000 ms, respectively. Dashed lines (color-coded) indicate average population firing rates. Vertical black lines flank the sample (500 – 1000 ms) and test (2500 – 3000 ms) stimulus intervals. Horizontal black line marks chance level decoding accuracy. (c) Decoding accuracy of the sample stimuli. Similar to (b) but impulse stimuli were applied at systematically varied delays (coloured bars). The colours of the traces correspond to the different flashes and highlight the changes in decoding accuracy induced by the flashes. Color-coded dashed lines indicate average population firing rates.

To investigate the time course of these impulse effects at higher temporal resolution, we systematically varied the timing of the impulse, increased the delay interval from 1500 ms to 1800 ms, reduced the duration of the sample stimulus from 500 to 400 ms and used only full screen white flashes of 100 ms duration (Figure 2c). This allowed us to assess the impulse effects with a temporal resolution of 200 ms across a total of eight experimental sessions. Again, the impulse stimuli led to a transient enhancement of decodability of the sample stimulus, and this enhancement of decodability was closely related in time with the transient increase in firing rate (Figure 2c). This impulse-related increase in decoding performance was also observed in the passive viewing experiments (Supplementary figure 7). These results suggest that the trailing trace of sample stimulus-specific information can be transiently enhanced as the V1 neurons are driven to fire by an unspecific impulse stimulus.

Enhanced stimulus trace with increased neuronal firing could simply be due to the fact that decodability of population vectors increases with discharge rate (Nikolić et al., 2009). However, this relationship does not always hold. In the DMS task (Supplementary figure 8) as well as its passive viewing version (Supplementary figure 9), stimuli evoked higher population firing rates when they were presented as sample rather than test. This difference was significant only in the last sustained response phase (300 – 350 ms after stimulus onset: *t* = 5.67, *p* = 0.00238, n = 6 sessions. *t*-test) but not during the response onset transient (e.g., 50 – 100 ms: *t* = 1.15, *p* = 0.303). However, the accuracy of decoding stimulus identity was higher for responses to the test than the sample stimulus, during both transient and sustained response phases (Supplementary figure 8 and Supplementary figure 9). Reduced firing rate and better decodability to the test stimulus also held when we performed the same analyses on nonmatch trials only (Supplementary figure 10), to rule out potential effects of repeated exposure to the same stimuli as is the case in the match trials. Thus, better decodability must have been due to other reasons than enhanced discharge rate.

### Reduced firing to stimuli matching priors

After the animal had learned the DMS task, we investigated whether the animal could learn implicit probabilistic associations between sample and test stimuli and use this information in the DMS task to predict the nature of the test stimulus given a particular sample. Predicted stimuli evoke smaller responses than unexpected stimuli (Alink et al., 2010; Peter et al., 2019; Uran et al., 2022). Therefore, we examined whether the same holds for test stimuli that were predicted with high or low probability by the sample stimuli. To this end, we modified the DMS task by introducing probabilistic pairing between sample and test stimuli in the nonmatch condition, such that the sample stimulus would predict with variable probability the upcoming test stimulus. Specifically, as shown in Figure 3a, in nonmatch trials (50% of all trials) the sample stimulus was followed by one of the two test stimuli with either high (40%, e.g., onion to kiwi, apple to paprika) or low (10%, e.g., onion to paprika, apple to kiwi) probability. The occurrence of sample-test pairs in the two probability conditions was counterbalanced. The other 50% of trials were match trials: each sample stimulus was followed by itself. In each session, we used two stimuli as sample and another two stimuli as test. The same set of four stimuli was used for the experimental sessions performed within a week to permit enough repetitions for the learning of the probabilistic association but different sets of stimuli were used for sessions in different weeks.

**Figure 3.**
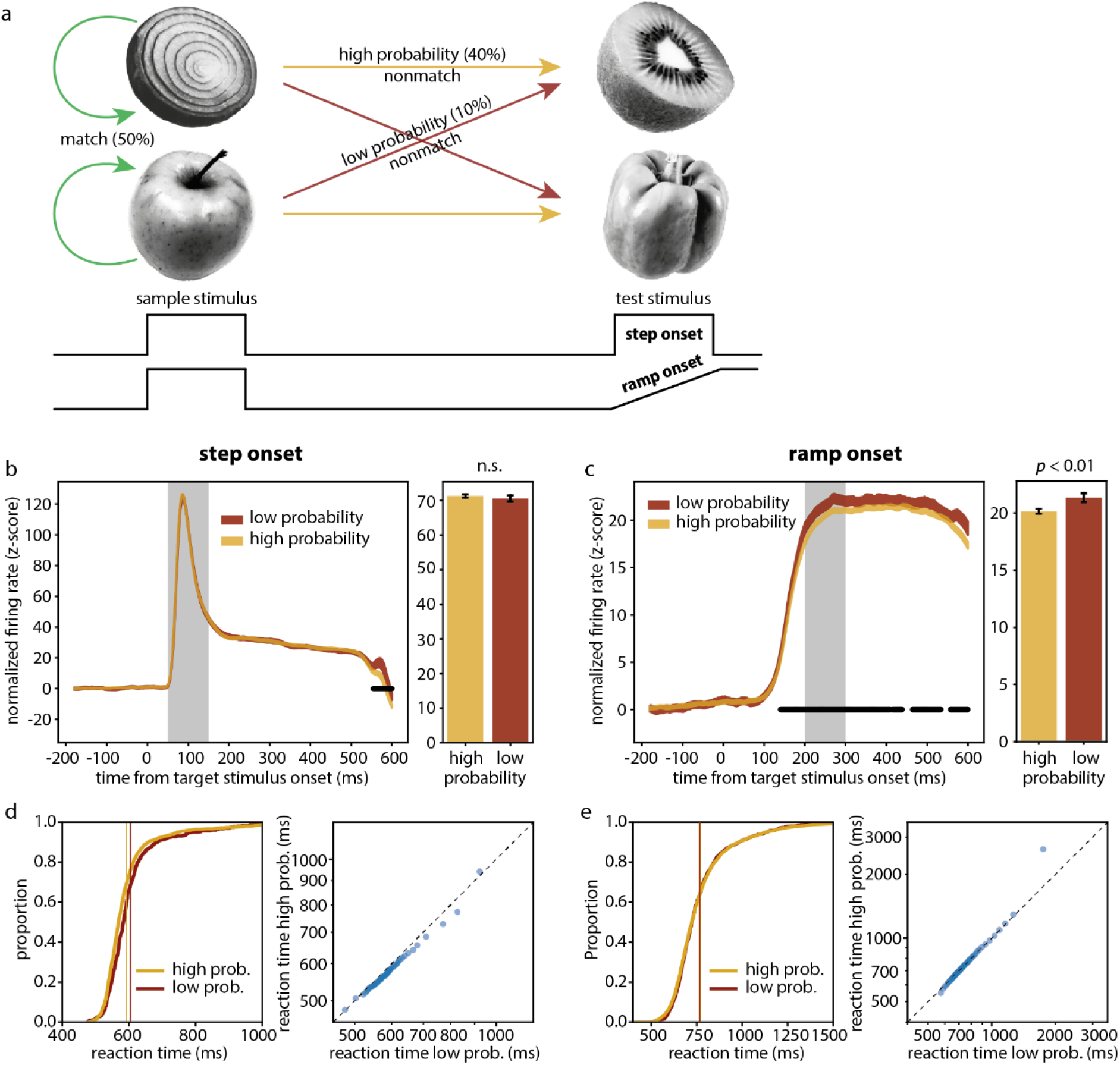
Neural and behavioural results for learning implicit probablistic associations. (a) Task structure. (b) Traces: normalized firing rate responses to high- and low-probability test stimuli in step onset condition. Bar plots: average firing rates (measurement interval marked in grey). Width of the traces and error bars represent 95% confidence interval. Black line marks statistically significant differences between high- and low-probability conditions. Note: the spurious difference at 550-600 ms is caused by the fact that trials are cut off at behavioural response time and that response time is shorter for high-probability condition. (c) Same convention as (b) but for ramp onset condition. (d) Left: cumulative density functions of reaction times in step onset condition. Vertical lines mark average reaction time. Right: quantile-quantile plot between reaction times for low (abscissa) and high (ordinate) probability conditions. Diagonal dashed line marks identity. (e) Same convention as (d) but for ramp onset condition.

To test whether neuronal responses to the test stimulus differed in high vs. low probability conditions, we computed test-evoked population firing rates (z-scored to pre-sample baseline per channel, summed across channels and normalized for pooling of trials from different sessions). We found no difference between high and low probability conditions (Figure 3b. Average firing rate within 50 – 150 ms after test onset: 71.31 ± 0.24 (s.e.m.) for high probability, 70.62 ± 0.47 for low probability; *t* = 1.28, *p* = 0.20). One reason for the lack of differences could be that subtle effects might have been overridden by the sharp initial transient responses caused by sudden stimulus onset. To examine this possibility, we slowly ramped up the intensity of the test stimulus (Figure 3a), expecting that such gradual visual input would dampen the abrupt increase in firing rate. The duration of the ramp was kept fixed for each session, but we varied this parameter across sessions (Monkey 1: 500 ms, 8 sessions; 2000 ms, 4 sessions; 3000 ms, 8 sessions. Monkey 2: 2000 ms, 14 sessions). Interestingly, in the experiments with ramping stimuli, for Monkey 1 the firing rate responses were weaker (*t* = –4.84, *p* = 1.34 × 10^-6^, all ramp durations combined, n = 20) to the test stimuli associated with high (20.15 ± 0.10 s.e.m., averaged within 200 – 300 ms after test onset) than low (21.33 ± 0.20) probability (Figure 3c). This difference was already evident in the early response phase (Figure 3c). As sanity check we compared the firing rates evoked by the same stimuli when they were in the sample position. There were no differences in high and low probability conditions (Supplementary figure 11a). To examine whether this probability-dependent difference in firing rate to the test stimulus was related to learning, we stratified the data into early (first experience with each set of stimuli and their associations) and late (last experience with the same sets of stimuli) sessions (200 trials in each of the early and late sessions, respectively, to equalize sample size), and found that the reduction of responses to high-probability test stimuli was only present in the late sessions (Supplementary figure 11c-d, 22.43 ± 0.33 for high probability, 24.078 ± 0.64 for low probability, *t* = –2.08, *p* = 0.038) and not in the early sessions (19.93 ± 0.43 for high probability, 21.31 ± 0.87 for low probability, *t* = –1.31, *p* = 0.19). The probability-dependent response difference in late rather than early sessions suggest the possibility that this effect was due to learning. However, we were unable to reproduce this result in Monkey 2 (Supplementary figure 12a).

To test whether probabilistic association between sample and test stimuli had an effect on the animal’s behaviour, we analysed the animal’s reaction (lever pressing) time after test stimulus appearance. For Monkey 1, when the test was presented with step onset, the reaction time was shorter for high (592.33 ± 98.97 ms, s.d.) than low (604.53 ± 96.15 ms) probability test stimuli (Figure 3d, *t* = –2.60, *p* = 9.29 × 10^-3^, *t*-test on log-transformed reaction time to ensure normality). However, when the test stimulus was presented with ramp onset, there was no difference in reaction time in the two probability conditions (Figure 3e, high probability 767.03 ± 178.87 ms, low probability 766.93 ± 165.30 ms; *t* = –0.51, *p* = 0.61, *t*-test). The same results held for the stratification in early (high probability 941.05 ± 266.52, low probability 936.35 ± 239.92; *t* = –0.028, *p* = 0.98) and late (high probability 775.56 ± 149.75, low probability 744.28 ± 105.91; *t* = 1.61, *p* = 0.11) sessions (Supplementary figure 11e-f). Between early and late sessions, the animal’s response accuracy (Supplementary figure 11b) increased from 72.0 ± 2.8 % (s.e.m.) to 94.8 ± 2.2 %, and reaction time decreased from 913.5 ± 269.7 ms (s.d.) to 713.8 ± 167.6 ms. For Monkey 2, we only used ramped test onset, and found that reaction times were slightly shorter for high (487.44 ± 121.04 ms) than low (495.51 ± 132.75 ms) probability tests (Supplementary figure 12b), but the difference was not statistically significant (*t* = –1.73, *p* = 0.084, *t*-test). However, if we first calculated the average reaction time per session for high and low probability conditions, respectively, and then performed pair-wise statistics across sessions, Monkey 2 also seemed to respond faster to high (488.31 ± 36.67 ms) than low (496.50 ± 37.93 ms) probability tests (Supplementary figure 12d, *t* = –2.72, *p* = 0.0176, paired *t*-test). This pairwise comparison revealed that the average reaction time per session was systematically shorter for the high probability test, although pooling data from all sessions did not uncover any statistically significant differences. Therefore, it seems that the animals may have learned and took advantage of the pairing between sample and test stimuli, and responded faster to high-probability test stimuli.

## Discussion

In this study, we trained monkeys to perform a working memory task and investigated the effect of both task-related and task-irrelevant factors on V1 neural activity. We found that V1 neurons did not show persistent spiking activity during the memory retention interval. However, decoding analysis revealed that the population vector of spiking activity contained a robust trace of the information about the sample stimulus despite the low firing rate. This trailing memory trace was apparently not caused by the specific demands of the WM task because it was also present in the passive viewing tasks. The trace of the preceding sample stimulus could be reactivated by an impulse stimulus that was unrelated to the sample but enhanced firing rates. Moreover, we found in the second series of experiments, that the amplitude of responses to the test stimulus depended on its expected probability and was reduced when the test matched prior expectation.

Before discussing these findings a few methodological considerations are warranted. To examine the role of V1 in working memory, we as well as other investigators relied on decoding methods to extract working memory content from V1 activity. However, decodability of WM content from V1 activity does *per se* not imply an involvement of V1 in WM. A better strategy would be to examine whether decoders could differentiate between the items that are required to be retained in WM (e.g., “test”) and those that are cued to be ignored (“distractor”). Such a distractor design is common in human psychophysics but challenging for non-human primates. But even then, improved decodability for WM items could reflect top-down effects related to predictive coding or feature-specific attention rather than an involvement in the maintenance of WM contents. These problems could in principle be overcome with loss of function experiments, i.e. transient inactivation of V1 during the retention interval (Rademaker et al., 2017). Another problem is that we were not able to determine whether the lingering memory traces in the DMS task and the passive viewing control had the same format or structure. This question could be resolved by performing transfer decoding to examine whether decoders trained on the DMS task could generalize to the passive viewing task and *vice versa*. Unfortunately, this was not possible because we used different sets of stimuli across sessions and the recorded signals likely drifted over days. Furthermore, the response modifications associated with test probability were not replicable in the second animal. Possible reasons are fewer sessions and less ramp variations for Monkey 2 than Monkey 1, different eccentricity of recording sites (RFs) and sampling bias. However, the results from Monkey 1 were robust and the stratification test provided unequivocal evidence for a learning-dependent process. Finally, our failure to retrieve WM-related information from V1 spiking activity may have been due to limitations of the decoder. We used a linear decoder which may have missed information contained in higher-order correlations (Bányai et al., 2019) or the temporal order of responses (Yiling et al., 2023).

Despite the low activity during the delay interval, the population firing rate vector contained information specific for the sample stimulus. The fact that this information was present in both DMS and passive viewing tasks makes it unlikely that it is related to an intentional effort to remember the sample. It is also unlikely that the lingering stimulus trace reflects a retinal afterimage, because stimulus contrast was low. Moreover, grating stimuli, which are typically used to induce afterimages, did not produce such lingering stimulus traces. The fact that the weaker traces after gratings were associated with worse behavioural performance compared to conditions with natural stimuli might be taken as evidence that lingering V1 activity is actually involved in the maintenance of WM. However, the results of the passive viewing task obtained in a naive animal do not support this assumption. Therefore, we favour the interpretation that the lingering trace reflects a form of fading memory that is maintained by the intrinsic dynamics of recurrent cortical networks (Buonomano and Maass, 2009; Nikolić et al., 2009). If so, this raises the question why information about complex natural stimuli persists longer than information about the orientation of gratings. Our results let it appear unlikely that this is simply due to a prolongation of reverberating responses to natural stimuli. Another possibility is that natural scene stimuli engage a larger network of recurrently coupled visual areas than gratings because they also match priors stored in the functional architecture of higher cortical areas (Bányai et al., 2019). This would imply that cooperation among multiple visual areas can enhance the persistence of stimulus-specific correlation structures in V1 activity. This interpretation is supported by the recent observation that stimulus-specific information persists longer in population responses recorded from areas V1 and V4 if responses are evoked by natural scene stimuli rather than by manipulated stimuli in which certain statistical regularities of natural stimuli were removed (Yiling et al., 2023). Also in these experiments there was no simple relation between discharge rates and decodability. Taken together, this suggests that fading memory is not a simple consequence of prolonged reverberation, a possibility worth further examination.

One possibility is that information is stored in stimulus-specific synaptic modifications that persist without requiring any sustained activity (Erickson et al., 2010; Mongillo et al., 2008). The finding that intervening impulse stimuli that transiently increased firing rates enhanced decodability would agree with this interpretation. In simulations, latent synaptic memory traces could be reactivated by unspecific network-wide stimulation (Mongillo et al., 2008). Likewise, in experiments with human subjects, transcranial magnetic stimulation (TMS) or strong visual stimulation revived latent contents of working memory (Rose et al., 2016; Wolff et al., 2017). In our study, this reactivation of stimulus-specific response vectors was similar in the WM tasks and the passive viewing controls. This suggests that the mechanism responsible for the fading memory trace can be activated by passive exposure and does not involve attention. This is in line with results of experiments on fading memory performed under anaesthesia (Nikolić et al., 2009). However, this does not imply that the lingering trace cannot be exploited by WM when required, nor does it exclude an influence of WM content on V1 processes.

The stimuli appearing in the test position evoked lower firing rates but were more decodable than in the sample position, in contrast to previous reports where higher firing rates improved decodability (Nikolić et al., 2009). The present results bear similarities with bottom-up mechanisms, such as adaptation and repetition suppression. However, several observations render classical repetition suppression unlikely. For briefly presented stimuli (400 to 500 ms) as used for our sample stimuli, adaptation acts mainly on the early transient response component, has only a weak or no influence on the late response phase (Liu et al., 2009; Muller et al., 1999; Patterson et al., 2013; Priebe et al., 2002) and vanishes within a few hundred milliseconds (Cohen-Kashi Malina et al., 2013; Patterson et al., 2013; Priebe et al., 2002). By contrast, in our experiments, only the late response phase was attenuated. Moreover, the effects were the same in nonmatch trials where the test stimulus was preceded by a different, therefore non-adapting, sample stimulus. Repetition suppression can also be excluded because its manifestation requires repeated exposure over minutes or hours. Another reason for the attenuation of responses to the test stimulus could be the predictability of the time of appearance and/or the need to respond to it. Since the trial timing was fixed, the monkey could predict precisely when the test stimulus would appear. Expectation and predictability have been shown to reduce neuronal firing (Alink et al., 2010; Meyer and Olson, 2011; Parras et al., 2017; Schwiedrzik and Freiwald, 2017; Todorovic and de Lange, 2012; Wacongne et al., 2011), to improve stimulus encoding (Bell et al., 2016; Kok et al., 2012), and to enhance gamma synchronization (Engel et al., 2001; Lima et al., 2011; Vinck and Bosman, 2016) in sensory areas. Notably, Todorovic and de Lange (2012) showed that, in line with our results, expectation-dependent suppression influenced the late response component (100 – 200 ms) whereas repetition suppression acted on the early component (40 – 60 ms). In sum, we favour the interpretation that the differences in sample-vs. test-evoked responses result from temporal expectation.

The effect of expectation on V1 activity is also evident in the DMS task in which we varied the probability with which a sample stimulus predicted a particular test stimulus. The probabilistic association between sample-test pairs was designed to establish an internal prior which rendered the sample stimulus predictive of the upcoming test stimulus. We found that test stimuli of high-probability evoked reduced firing rates as compared to the low-probability stimuli. Because the same set of test stimuli was used in the two probability conditions and the stimulus set varied across sessions, test stimulus-specific differences in firing rate are unlikely to explain the effect. Moreover, the probability-dependent modification of firing rate emerged only in late but not in early sessions. As adaptation and repetition dependent effects are unlikely (see above), these observations suggest learnt predictability as a likely cause for the dependence of firing rate on the probability of the test stimulus. Although our data do not allow us to identify the site where the prior for the prediction is generated, the ramping paradigm revealed that the prior-associated effect kicked in already in the initial response phase, suggesting fast access to information about the nature of the expected stimulus. The reduction of responses to predictable stimuli is in agreement with reports of reduced activation of visual cortex by stimuli that comply with predictions (Alink et al., 2010; de Lange et al., 2018; Kok et al., 2012; Meyer and Olson, 2011; Peter et al., 2019; Schwiedrzik and Freiwald, 2017; Summerfield and de Lange, 2014; Uran et al., 2022). This suppression has been interpreted in the context of predictive coding as a mechanism to facilitate perceptual inference (Aitchison and Lengyel, 2017; Rao and Ballard, 1999; Singer, 2021; Singer and Lazar, 2016), to reduce redundant signals originating from compressible stimuli (Peter et al., 2019; Uran et al., 2022; Vinck and Bosman, 2016) and to improve stimulus representation (Bell et al., 2016; Kok et al., 2012).

In summary, the intrinsic dynamics of early visual areas are capable of maintaining re-activatable traces of complex visual stimuli. We propose as the most likely mechanism the fading memory that is characteristic of recurrent networks. These lingering memory traces do not seem to depend on active, attention-dependent WM processes but could of course support WM if required. Ample evidence indicates that the responses of V1 neurons depend to a large extent on the match between sensory evidence and priors. Some of these priors have been acquired during evolution, are complemented by experience-dependent developmental pruning and perceptual learning and are stored in the functional architecture of the visual cortex (reviewed in Singer (2021)). Other priors are derived from the actual context in which stimuli are presented (Lazar et al., 2021; Peter et al., 2019; Uran et al., 2022). Our results indicate that also predictions derived from learnt associations impact responses in V1. Although initial acquisition of these associations between temporally distant stimuli very likely involves WM, once established, these associations must be stored in long-term memory. As our results indicate, this covert knowledge about the likelihood of the appearance of a particular stimulus is available in primary visual cortex. We consider it unlikely that the association between sample and test is formed in V1 and therefore favour the interpretation that the priors set up dynamically by the sample stimulus are conveyed to V1 by top down projections.

## Methods

### Behavioural task

Results presented here were obtained from two adult rhesus monkeys (*Macacca mulatta*. Monkey 1: male, 11 kg, 12 years old. Monkey 2: female, 9 kg, 17 years old). All experimental procedures were in compliance with the German and European regulations on laboratory animal protection and welfare, and were approved by the local authority (Regierungspräsidium Darmstadt). The monkey was seated 60 cm in front of a screen (Samsung SyncMaster 2233RZ; 120 Hz refresh rate) inside a dark booth. The monkey initiated a trial by fixating at a white fixation dot displayed at the centre of the screen, and had to maintain fixation on the fixation dot until the trial ended. The eye position was monitored with the EyeLink tracker (SR Research, Ottawa, Ontario, Canada).

In the delayed match-to-sample (DMS) task, two stimuli were presented sequentially and the monkey had to report whether the two stimuli were the same (match) or different (nonmatch). A trial started with a fixation period of 500 ms, during which the screen was blank. Then the first stimulus (sample) was presented for 500 ms, followed by a delay period of 1500 ms after which the second stimulus (test) was presented. The test stimulus disappeared once the monkey responded and kept on for maximally 500 ms in case of delayed responses. The monkey had to respond by moving a two-way mechanical lever; forward in match trials and backwards in nonmatch trials. Monkeys were not rewarded for responding swiftly. The number of match and nonmatch trials was pseudo-randomized to be equal. A correct response was rewarded with a drop of water or juice. If the monkey broke fixation (1.3° around fixation point), or moved the lever before test onset, the trial was aborted.

The trial structure of the passive viewing control experiment mimicked that of the DMS task. In the passive viewing task, the monkey had to maintain fixation and respond to a colour change of the fixation spot in order to be rewarded. During fixation, the stimuli were presented as in the DMS task, but were irrelevant to the animal. The fixation dot changed its colour to either green or blue, requiring forward or backward moves of the lever, respectively.

### Stimulus design

The stimuli were images of single isolated objects with transparent background. The visible region of the image was normalized to equal pixel intensity (0.5) and root-mean-square contrast (0.275), and covered the classical receptive fields of the recorded multi-units. The image size was 160 × 160 pixels (4.4° visual angle) for Monkey 1, and 250 × 250 pixels (6.9° visual angle) for Monkey 2. The images were shown at 50% transparency (alpha = 0.5) to reduce the potential influence of visual adaptation. The background colour of the display screen was 0.5 grey level throughout the experiment. In the standard DMS task, a set of three stimuli were used in each session. The set of images varied between sessions. Each image could appear in both the sample and test positions, at pseudo-randomized equal probability.

In the probabilistic DMS task, a set of four stimuli were used in each session, and the pairing between sample and test stimuli was additionally manipulated. Only two of the four stimuli could appear as sample (and as test in match trials), and the other two stimuli only appeared as test stimuli (i.e., only appeared in nonmatch trials). In nonmatch trials (50% of all trials; the other 50% are match trials), the sample stimulus was followed by one of the two test stimuli with either high (40%) or low (10%) probability. The occurrence of sample-test pairs in the two probability conditions was counterbalanced, such that in nonmatch trials both test stimuli appeared at equal probability and only their probabilistic pairing with the proceeding sample stimulus was shuffled.

### Electrophysiology

Both monkeys were chronically implanted in the left hemisphere over V1 with a microdrive system (Gray Matter Research, Bozeman, Montana, USA) which had 32 individually movable microelectrodes. Data acquisition was performed using the TDT system (Tucker-Davis Technologies, Alachua, Florida, USA). The signal was amplified and digitalized at 25 kHz (TDT PZ5 NeuroDigitizer). This raw signal was bandpass-filtered between 300 – 4000 Hz to extract multi-unit spiking activity (MUA), and low pass-filtered at 300 Hz and down-sampled with a decimation factor of 24 to about 1 kHz to retrieve the local field potential (LFP). MUA was isolated using the online detection algorithm in the OpenEx software (Tucker-Davis Technologies). Events crossing a threshold of 4 times the standard deviation of the filtered spiking band activity were considered as spikes and analysed further.

### Data analysis

All decoding analyses were based on linear discriminant analysis (LDA) classifiers. Firing rates across channels were treated as independent variables, i.e. predictors, with repeated measurements across trials. To test for stimulus-specificity, image identity was used as class label. For time-resolved decoding, independent classifiers were trained at successive time points. In most decoding analyses, firing rate was calculated by binning spikes in moving windows of 100 ms and steps of 50 ms. For finer timescale comparison of decoding sample-vs. test-evoked activity, we used smaller windows (50 ms) and step sizes (10 ms). Decoding accuracy for each session was measured by averaging the cross-validated classification performance over 20 repeated stratified sampling of the dataset (20 folds). Decoding accuracy values were averaged over sessions.

## Supplementary Figures

**Supplementary figure 1.**
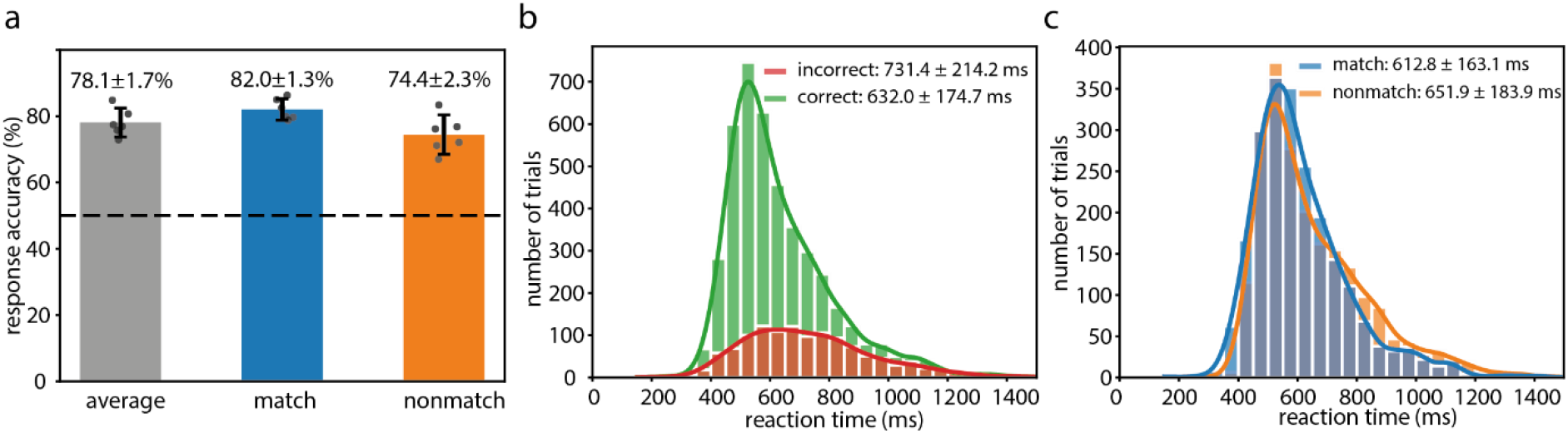
Behavioural performance in the DMS task (Monkey 1). (a) Response accuracy for all trials (grey), match trials (blue) and nonmatch trials (orange). Error bars denote 95% confidence intervals. Error numbers in the notations above the bar plots represent standard error of the mean. (b) Reaction time distribution for correct and incorrect trials. Error numbers in the legend denote standard deviation. (c) Reaction time distribution for match and nonmatch correct trials. Error numbers in the legend denote standard deviation.

**Supplementary figure 2.**
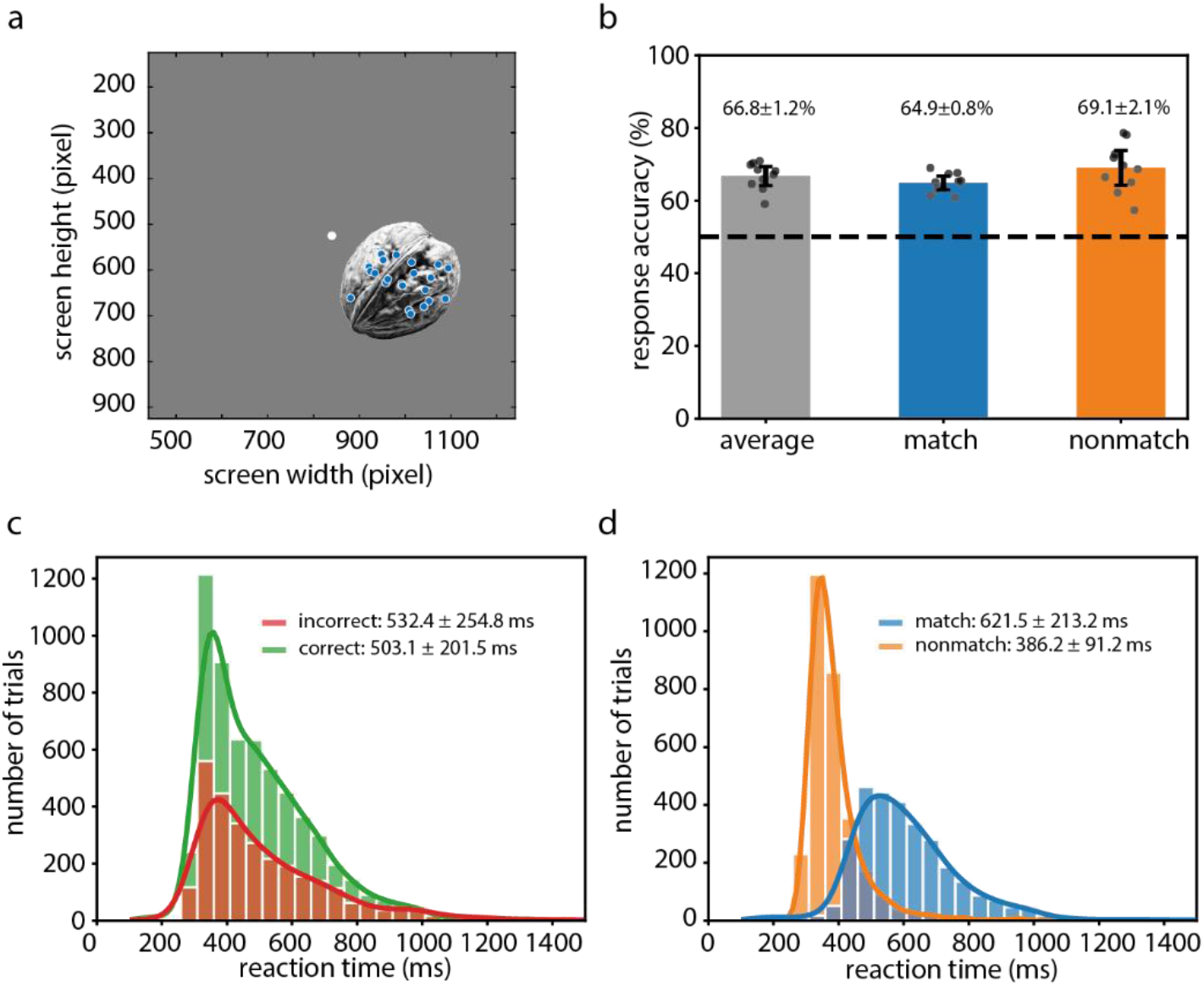
Example stimulus, RF locations and behavioural performance of Monkey 2 in the DMS task. (a) Stimulus position on the screen and receptive fields (blue dots). Specifically, stimuli subtended 7.84° of visual angle (DVA). Their centre was 4.05° lateral to the vertical and 2.70° below the horizontal meridian. (b-d) Same convention as in Supplementary figure 1a-c. Behavioural performance was assessed from 10 sessions (n=10).

**Supplementary figure 3.**
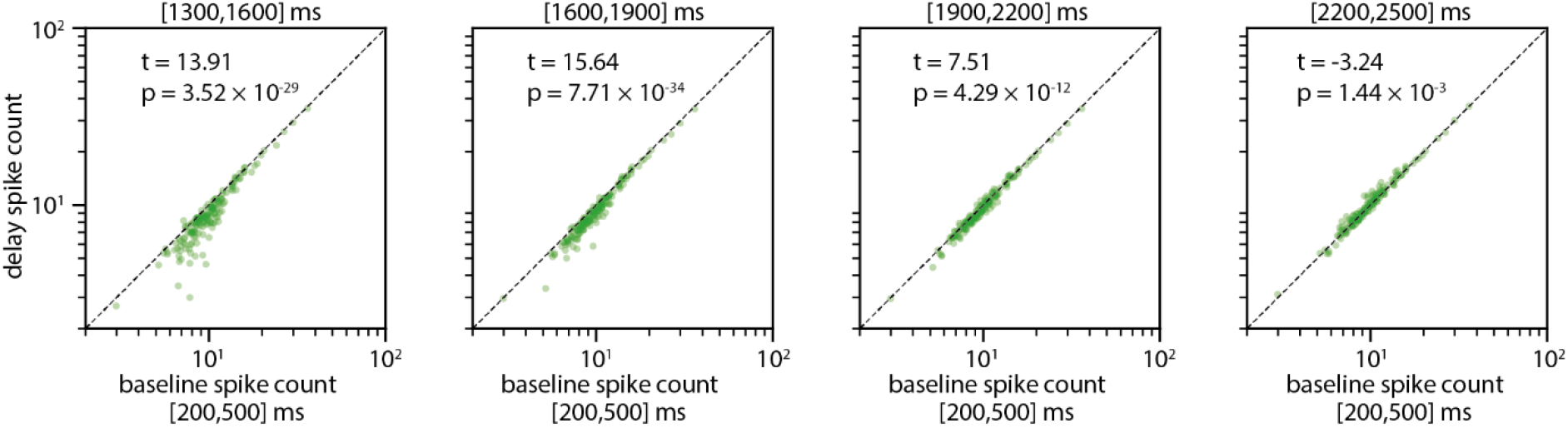
Comparison of spike counts between delay period and pre-stimulus baseline. Spike counts measured in different stages of delay interval (y-axis; 300ms time windows, marked in the titles of panels) plotted against baseline spike counts (x-axis; equal time window). Dotted lines indicate equality.

**Supplementary figure 4.**
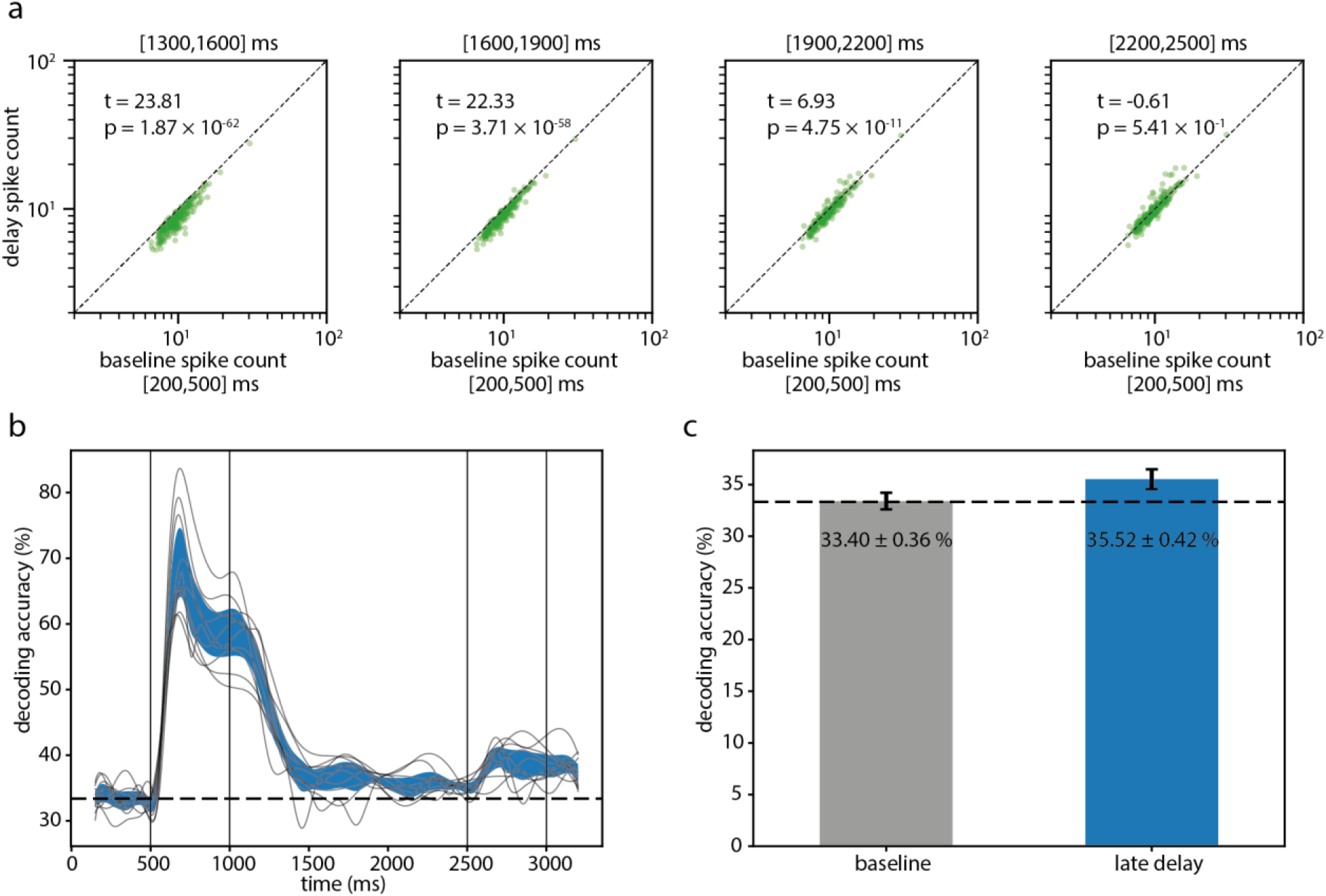
Robust trace of sample stimulus information during the delay interval (results from Monkey 2). (a) Same convention as Supplementary figure 3. (b-c) Same conventions as Figure 1d-e of the main text. Baseline: t = 0.18, p = 0.861. Delay: t = 4.88, p = 0.000872. Between: t = −3.82, p = 0.00408.

**Supplementary figure 5.**
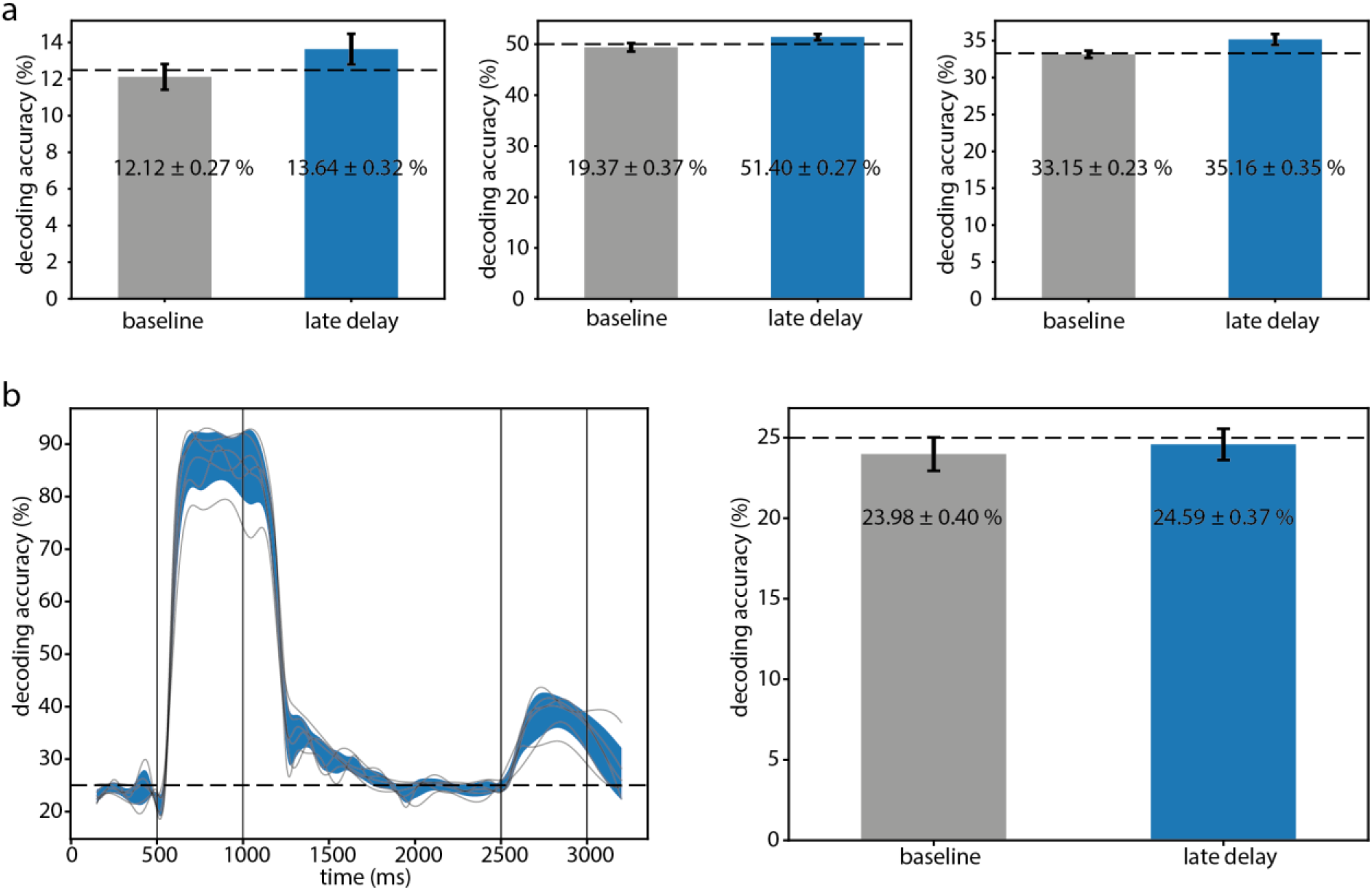
Robust trace of stimulus information during the delay period. Summary of results obtained from a few other independent DMS experiments (Monkey 1). (a) The accuracies of decoding sample stimulus identity for the baseline period (grey) and the last 500 ms of the delay interval (blue). The three panels represent three sets of experiments that used 8 (left, n = 6 sessions), 2 (middle, n = 12 sessions), and 3 (right, n = 22 sessions) stimuli per session, respectively. Note that the stimuli varied for both different sessions and experiments. In all three experiments, the decoding accuracies for the sample stimuli in the late delay period were significantly above chance level and higher than baseline level. All error bars denote the 95% confidence interval. (b) A fourth DMS experiment that used four grating stimuli per session (n = 6 sessions). The grating orientations varied between sessions. Left: Time resolved decoding accuracy for the sample stimuli. Right: Same arrangement as in (a). Note that in this experiment, the decoding accuracy for the sample stimuli in the late delay interval was not different from the chance level (25%, t = −1.00, p = 0.362) or the baseline level (t = −0.99, p = 0.366).

**Supplementary figure 6.**
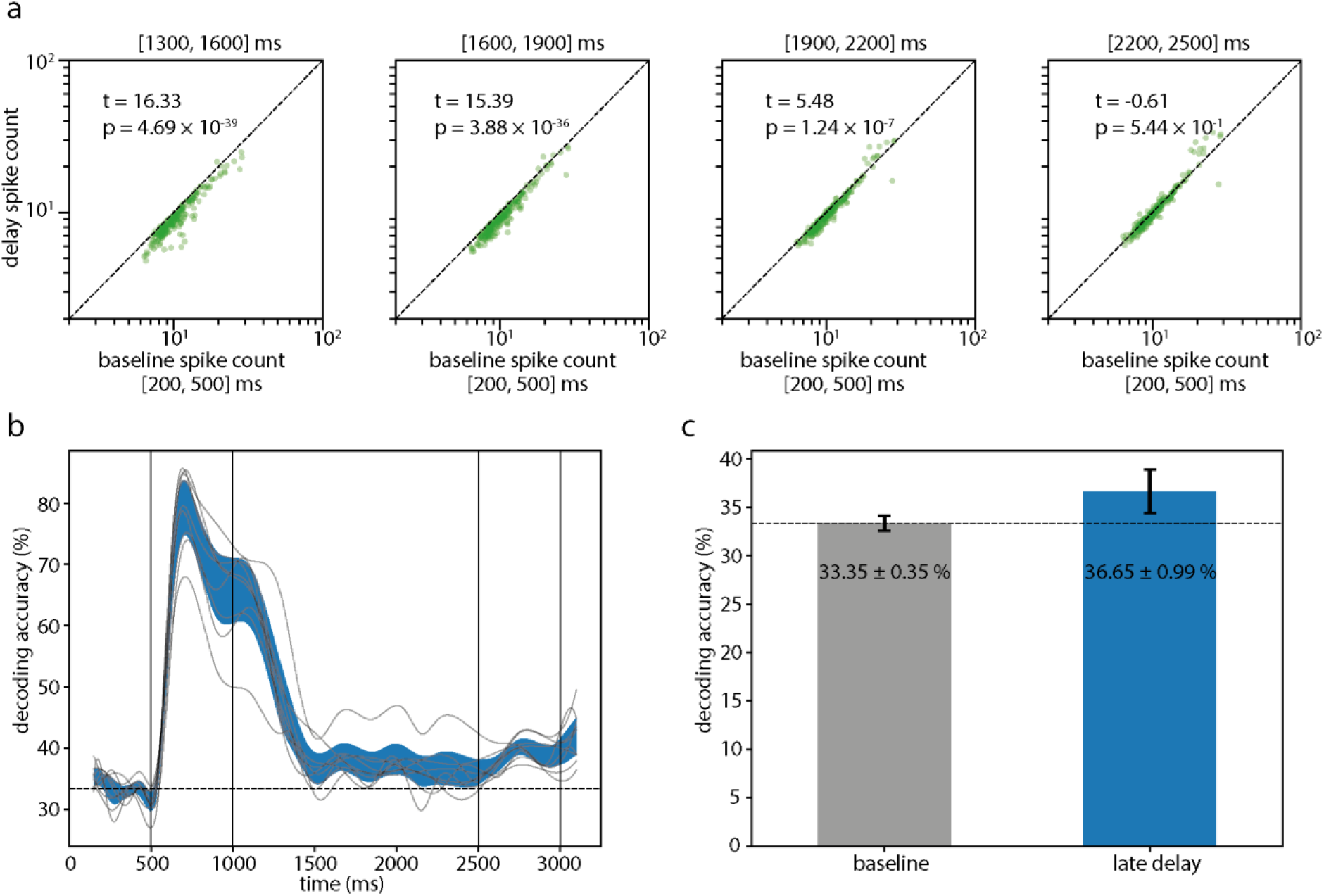
Persistence of sample stimulus information during the delay period of the passive viewing control experiment. (c) Average decoding accuracy during the late delay interval was significantly above chance level (t = 3.17, p = 0.0132, n = 9 sessions, t-Test) and above baseline level (t = −3.13, p = 0.141). The decoding accuracy during the baseline period was not significantly different from chance level (t = 0.06, p = 0.957).

**Supplementary figure 7.**
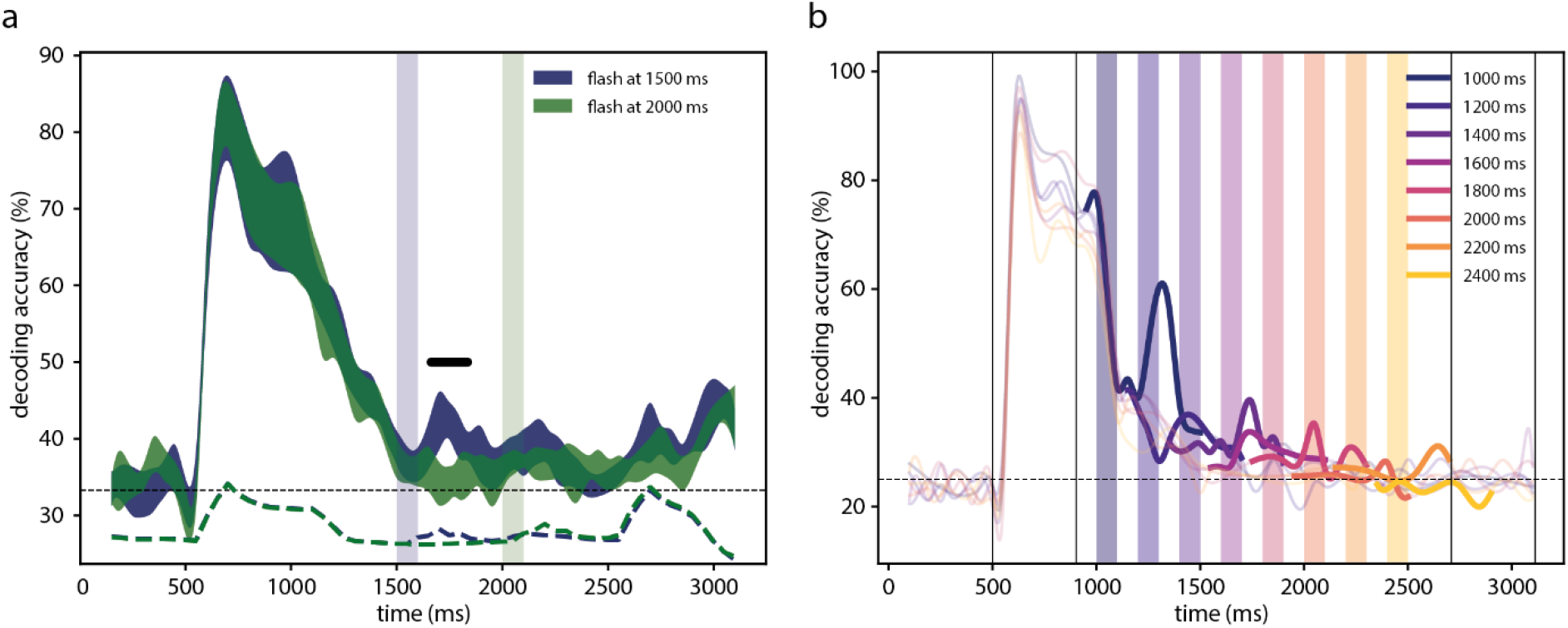
Visual stimulation reactivates latent stimulus trace. (a) Results from the DMS experiment in Monkey 2. (b) Results from passive viewing experiments (Monkey 1). Same conventions as for Figure 2 in the main text.

**Supplementary figure 8.**
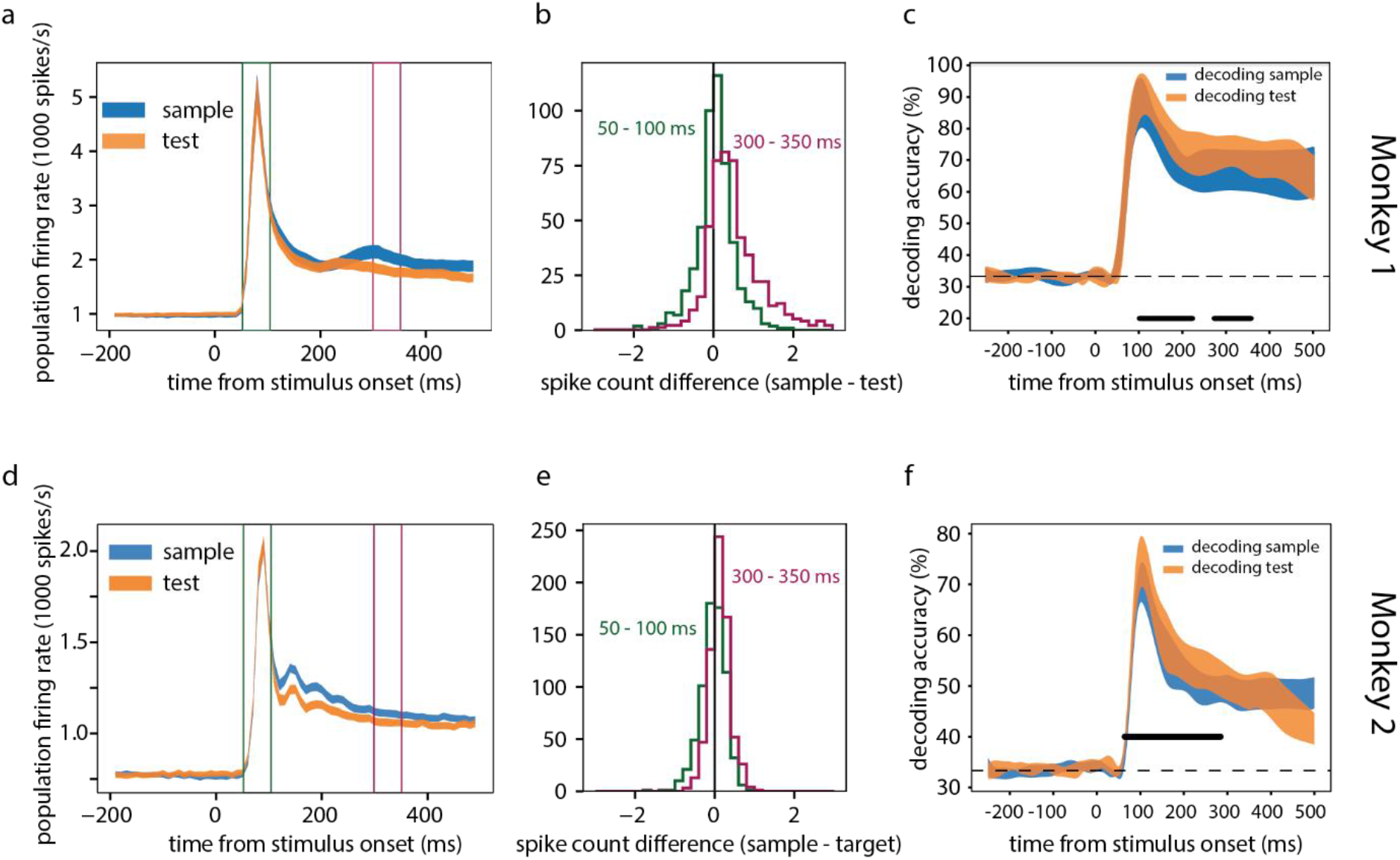
Lower firing rate and higher decodability to test than sample stimuli. (a) Population firing rate (summed across all multi-units) evoked by stimuli in the sample and test position, respectively. (b) Difference in evoked spike count (sample – test, per multi-unit channel) during transient (50 – 100 ms) and late response phase (300 – 350 ms). See time windows marked in (a). (c) Accuracy of decoding sample and test stimulus identity. Shades denote 95% confidence intervals. Black bars mark regions of statistical difference. (d-f) Same convention as in (a-c) but for Monkey 2. Difference in the transient phase (50 – 100 ms): t = −1.65, p = 0.133, n = 10 sessions. Difference in the late phase (300 – 350 ms): t = 8.43, p = 1.45 × 10^-5^.

**Supplementary figure 9.**
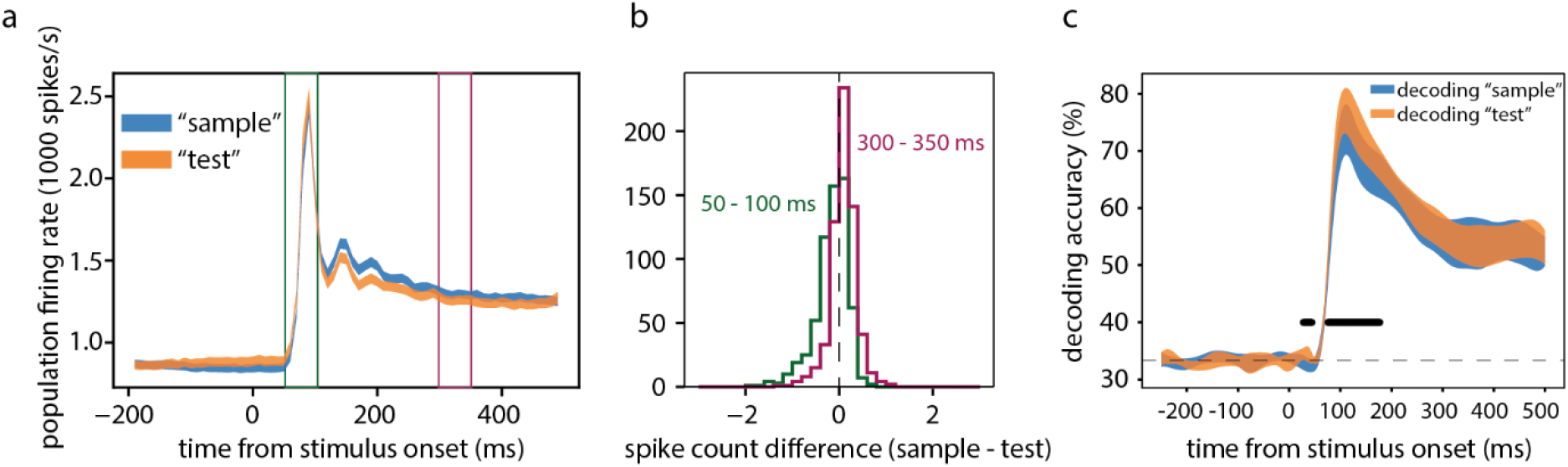
Lower firing rate and higher decodability to “test” than “sample” stimuli. Results from the passive viewing control version of the DMS task. Same convention as in Supplementary figure 8. Firing rate difference in the transient phase (50 – 100 ms): t = −4.00, p = 3.94 × 10^−3^, n = 9 sessions. Difference in the late phase (300 – 350 ms): t = 5.13, p = 8.93 × 10^−4^.

**Supplementary figure 10.**
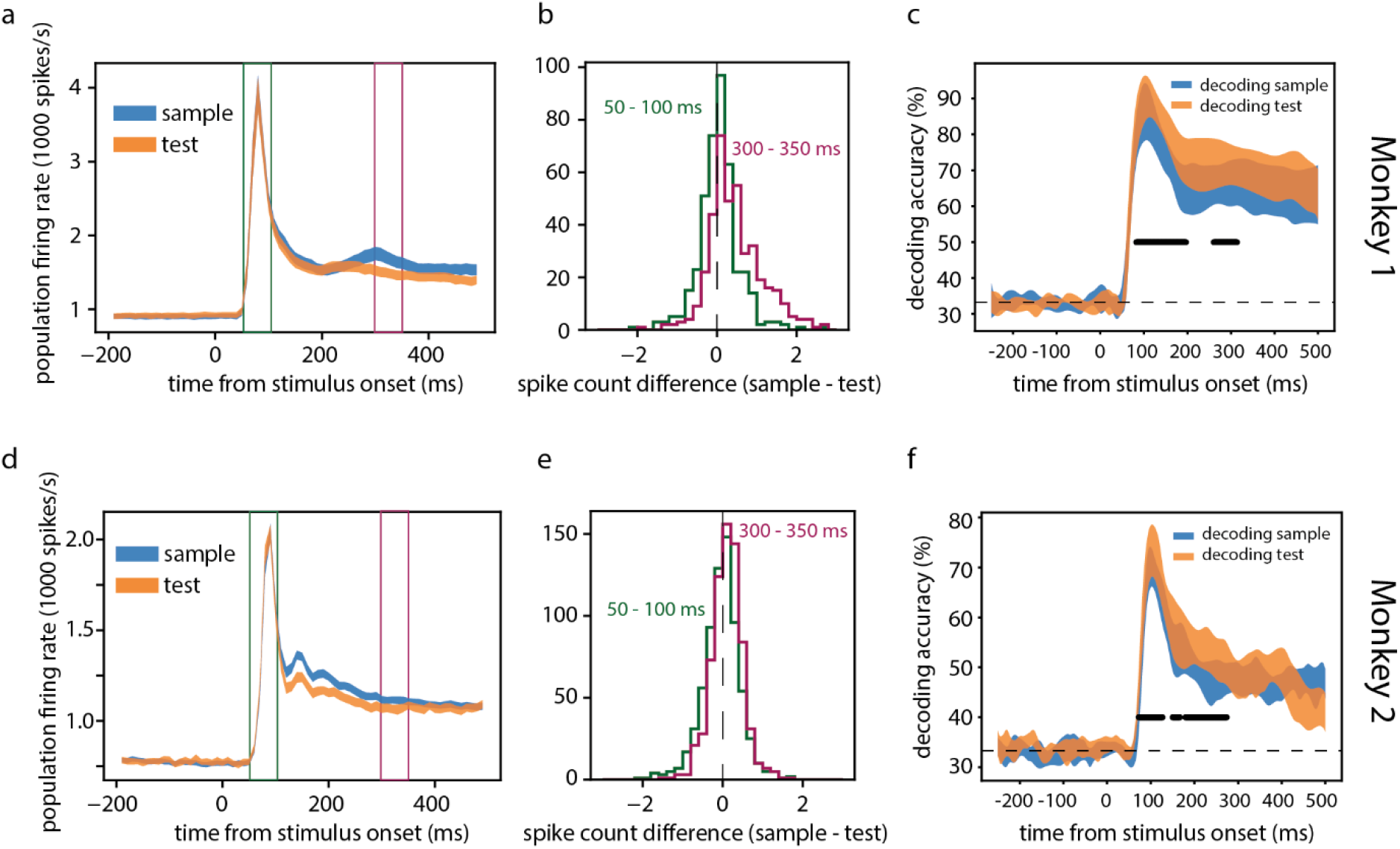
Lower firing rate and higher decodability to test than sample stimuli (nonmatch trials only). Same convention as in Supplementary figure 8.

**Supplementary figure 11.**
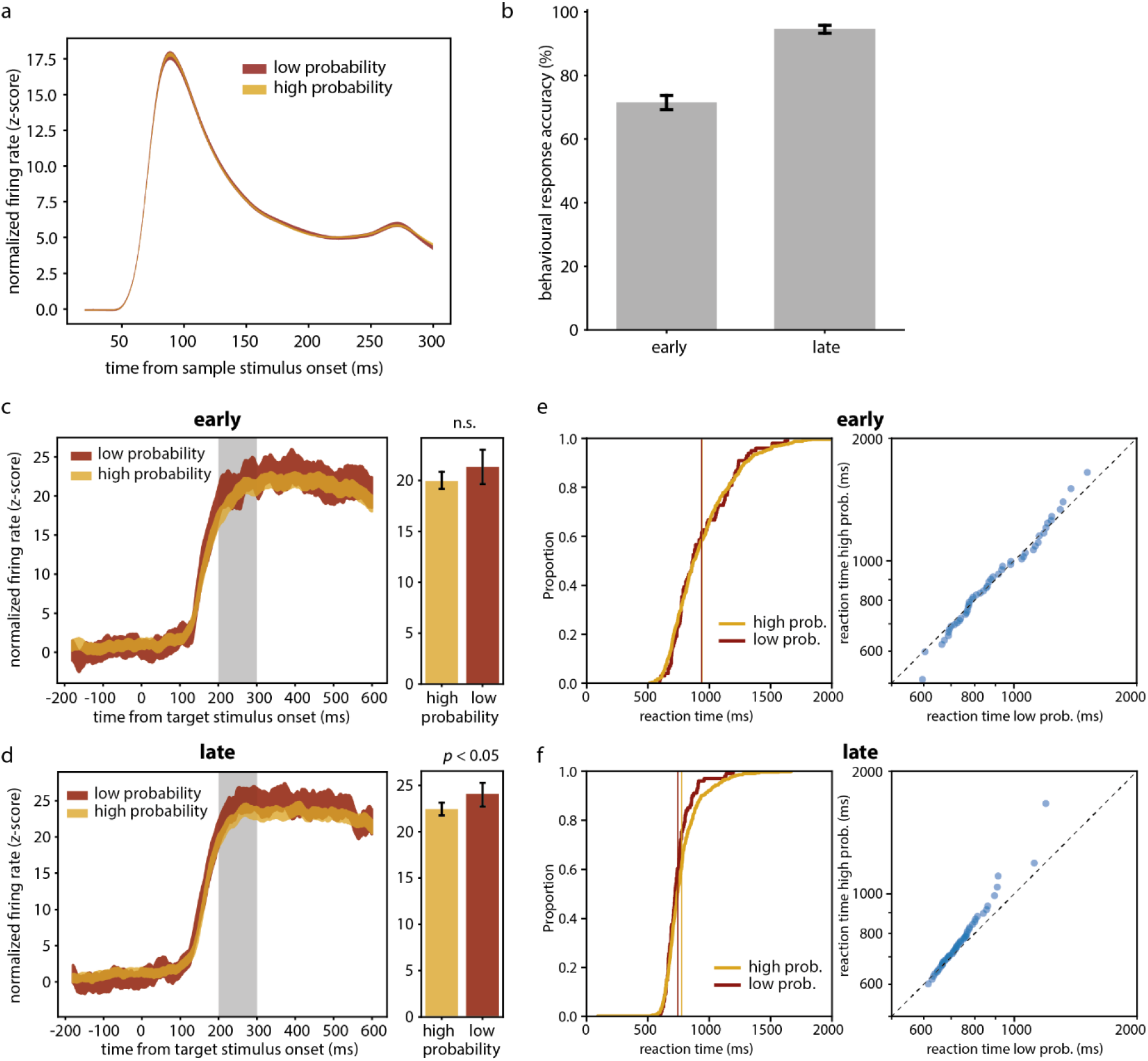
Comparison between early and late sessions of learning associations between stimuli (Monkey 1). (a) Normalized firing rate responses to sample stimuli in low vs. high probability conditions. (b) Behavioural response accuracy between early and late sessions. (c) & (d) Same convention as Figure 3c but separated for early (c) and late (d) sessions. (e) & (f) Same convention as Figure 3e.

**Supplementary figure 12.**
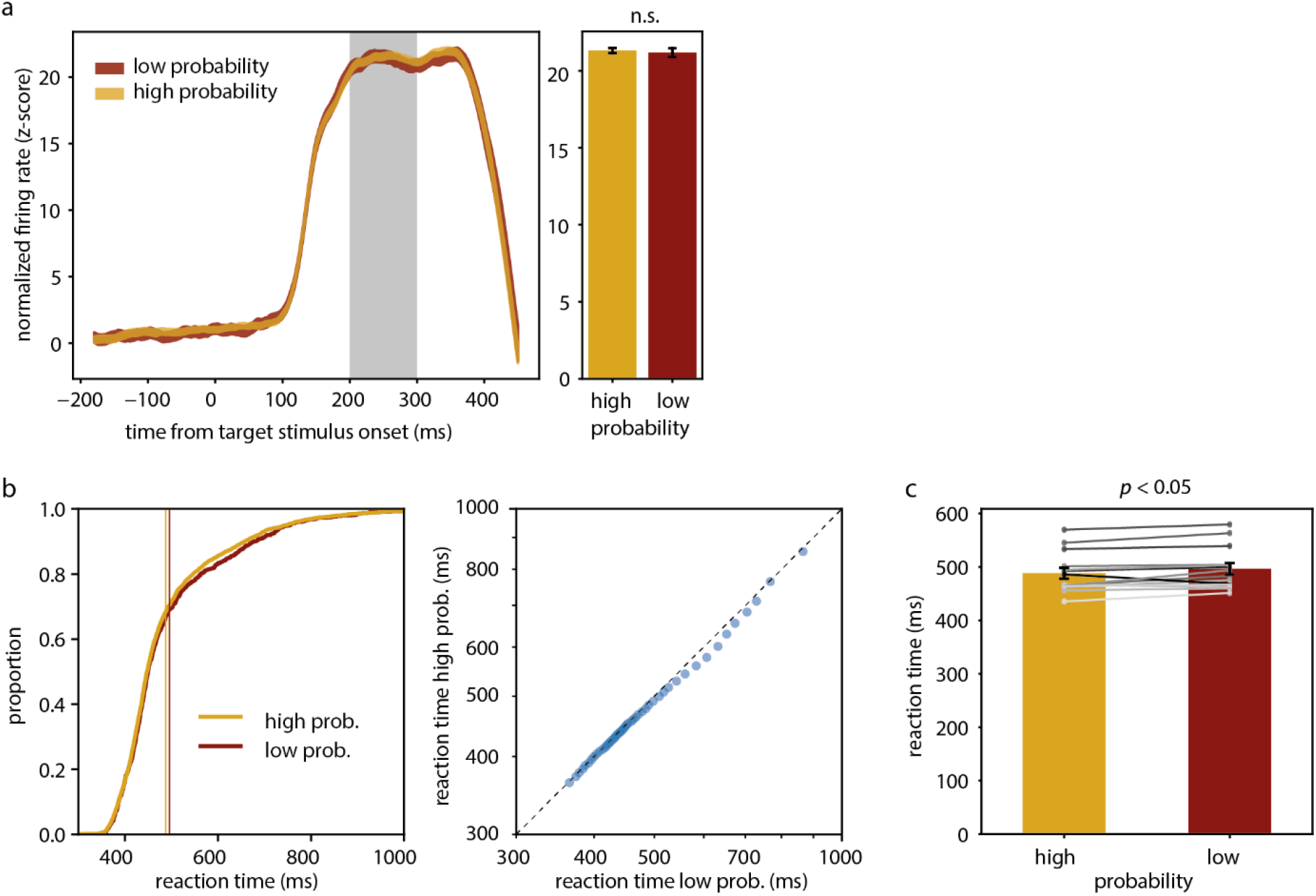
Neural and behavioural results for learning implicit probablistic association. Results from Monkey 2. (a) Same conventions as Figure 3c. (b) Same conventions as Figure 3e. (c) Reaction time in high and low probability conditions. Connected dots denote average reaction per session. Grey level marks the order of sessions (dark to light grey: early to late sessions).

## Acknowledgements

We thank the Ernst Strüngmann Foundation, Max Planck Society, the Human Frontier Science Program (HFSP RGP0044/2018) and the Deutsche Forschungsgemeinschaft (DFG Reinhart Koselleck Project 325248489) for supporting this work. We are grateful to Rosanne Rademaker and Michael Wolff for sharing helpful comments on the manuscript.

## Author Contributions

Experimental design: YYL, KS, AL, WS.

Experimentation: YYL, JKL, KS.

Data analysis: YYL.

Writing: YYL and WS.

Supervision: WS.

## Competing Interests

The authors declare no competing interests.

## Notes

### Competing Interest Statement

The authors have declared no competing interest.

## References

Aitchison, L., and Lengyel, M. (2017). With or without you: predictive coding and Bayesian inference in the brain. Current opinion in neurobiology 46, 219–227.

Alink, A., Schwiedrzik, C.M., Kohler, A., Singer, W., and Muckli, L. (2010). Stimulus predictability reduces responses in primary visual cortex. J Neurosci 30, 2960–2966.

Bányai, M., Lazar, A., Klein, L., Klon-Lipok, J., Stippinger, M., Singer, W., and Orbán, G. (2019). Stimulus complexity shapes response correlations in primary visual cortex. Proceedings of the National Academy of Sciences of the United States of America 116, 2723–2732.

Bell, A.H., Summerfield, C., Morin, E.L., Malecek, N.J., and Ungerleider, L.G. (2016). Encoding of Stimulus Probability in Macaque Inferior Temporal Cortex. Curr Biol 26, 2280–2290.

Bergmann, J., Genc, E., Kohler, A., Singer, W., and Pearson, J. (2016). Neural Anatomy of Primary Visual Cortex Limits Visual Working Memory. Cereb Cortex 26, 43–50.

Brincat, S.L., Donoghue, J.A., Mahnke, M.K., Kornblith, S., Lundqvist, M., and Miller, E.K. (2021). Interhemispheric transfer of working memories. Neuron 109, 1055–1066 e1054.

Brodeur, M.B., Dionne-Dostie, E., Montreuil, T., and Lepage, M. (2010). The Bank of Standardized Stimuli (BOSS), a new set of 480 normative photos of objects to be used as visual stimuli in cognitive research. PLoS One 5, e10773.

Brodeur, M.B., Guerard, K., and Bouras, M. (2014). Bank of Standardized Stimuli (BOSS) phase II: 930 new normative photos. PLoS One 9, e106953.

Buonomano, D.V., and Maass, W. (2009). State-dependent computations: spatiotemporal processing in cortical networks. Nature Reviews Neuroscience 10, 113–125.

Christophel, T.B., Iamshchinina, P., Yan, C., Allefeld, C., and Haynes, J.D. (2018). Cortical specialization for attended versus unattended working memory. Nat Neurosci 21, 494–496.

Cohen-Kashi Malina, K., Jubran, M., Katz, Y., and Lampl, I. (2013). Imbalance between excitation and inhibition in the somatosensory cortex produces postadaptation facilitation. J Neurosci 33, 8463–8471.

Constantinidis, C., Funahashi, S., Lee, D., Murray, J.D., Qi, X.L., Wang, M., and Arnsten, A.F.T. (2018). Persistent Spiking Activity Underlies Working Memory. J Neurosci 38, 7020–7028.

D’Esposito, M. (2007). From cognitive to neural models of working memory. Philos Trans R Soc Lond B Biol Sci 362, 761–772.

D’Esposito, M., and Postle, B.R. (2015). The cognitive neuroscience of working memory. Annu Rev Psychol 66, 115–142.

de Lange, F.P., Heilbron, M., and Kok, P. (2018). How Do Expectations Shape Perception? Trends Cogn Sci 22, 764–779.

de Vries, I.E.J., Slagter, H.A., and Olivers, C.N.L. (2020). Oscillatory Control over Representational States in Working Memory. Trends Cogn Sci 24, 150–162.

Emrich, S.M., Riggall, A.C., Larocque, J.J., and Postle, B.R. (2013). Distributed patterns of activity in sensory cortex reflect the precision of multiple items maintained in visual short-term memory. J Neurosci 33, 6516–6523.

Engel, A.K., Fries, P., and Singer, W. (2001). Dynamic predictions: oscillations and synchrony in top-down processing. Nat Rev Neurosci 2, 704–716.

Erickson, M.A., Maramara, L.A., and Lisman, J. (2010). A single brief burst induces GluR1-dependent associative short-term potentiation: a potential mechanism for short-term memory. J Cogn Neurosci 22, 2530–2540.

Ester, E.F., Anderson, D.E., Serences, J.T., and Awh, E. (2013). A neural measure of precision in visual working memory. J Cogn Neurosci 25, 754–761.

Ester, E.F., Serences, J.T., and Awh, E. (2009). Spatially global representations in human primary visual cortex during working memory maintenance. J Neurosci 29, 15258–15265.

Fiebig, F., and Lansner, A. (2017). A Spiking Working Memory Model Based on Hebbian Short-Term Potentiation. J Neurosci 37, 83–96.

Funahashi, S., Bruce, C.J., and Goldman-Rakic, P.S. (1989). Mnemonic coding of visual space in the monkey’s dorsolateral prefrontal cortex. Journal of neurophysiology 61, 331–349.

Fuster, J.M., and Alexander, G.E. (1971). Neuron activity related to short-term memory. Science 173, 652–654.

Gray, C.M., Konig, P., Engel, A.K., and Singer, W. (1989). Oscillatory responses in cat visual cortex exhibit inter-columnar synchronization which reflects global stimulus properties. Nature 338, 334–337.

Gray, C.M., and Singer, W. (1989). Stimulus-specific neuronal oscillations in orientation columns of cat visual cortex. Proc Natl Acad Sci U S A 86, 1698–1702.

Haller, M., Case, J., Crone, N.E., Chang, E.F., King-Stephens, D., Laxer, K.D., Weber, P.B., Parvizi, J., Knight, R.T., and Shestyuk, A.Y. (2018). Persistent neuronal activity in human prefrontal cortex links perception and action. Nat Hum Behav 2, 80–91.

Harrison, S.A., and Tong, F. (2009). Decoding reveals the contents of visual working memory in early visual areas. Nature 458, 632.

Iamshchinina, P., Christophel, T.B., Gayet, S., and Rademaker, R.L. (2021a). Essential considerations for exploring visual working memory storage in the human brain. Visual Cognition 29, 425–436.

Iamshchinina, P., Christophel, T.B., Gayet, S., and Rademaker, R.L. (2021b). Understanding how analysis choices are essential for the meaningful interpretation of visual working memory data. Journal of Vision 21, 2721–2721.

Kaminski, J., Sullivan, S., Chung, J.M., Ross, I.B., Mamelak, A.N., and Rutishauser, U. (2017). Persistently active neurons in human medial frontal and medial temporal lobe support working memory. Nat Neurosci 20, 590–601.

Kapadia, M.K., Ito, M., Gilbert, C.D., and Westheimer, G. (1995). Improvement in visual sensitivity by changes in local context: parallel studies in human observers and in V1 of alert monkeys. Neuron 15, 843–856.

Kok, P., Jehee, J.F., and de Lange, F.P. (2012). Less is more: expectation sharpens representations in the primary visual cortex. Neuron 75, 265–270.

Kornblith, S., Quian Quiroga, R., Koch, C., Fried, I., and Mormann, F. (2017). Persistent Single-Neuron Activity during Working Memory in the Human Medial Temporal Lobe. Curr Biol 27, 1026–1032.

Kubota, K., and Niki, H. (1971). Prefrontal cortical unit activity and delayed alternation performance in monkeys. Journal of neurophysiology 34, 337–347.

Lara, A.H., and Wallis, J.D. (2015). The Role of Prefrontal Cortex in Working Memory: A Mini Review. Front Syst Neurosci 9, 173.

Lawrence, S.J.D., van Mourik, T., Kok, P., Koopmans, P.J., Norris, D.G., and de Lange, F.P. (2018). Laminar Organization of Working Memory Signals in Human Visual Cortex. Curr Biol 28, 3435–3440 e3434.

Lazar, A., Lewis, C., Fries, P., Singer, W., and Nikolic, D. (2021). Visual exposure enhances stimulus encoding and persistence in primary cortex. Proc Natl Acad Sci U S A 118.

Lima, B., Singer, W., and Neuenschwander, S. (2011). Gamma responses correlate with temporal expectation in monkey primary visual cortex. J Neurosci 31, 15919–15931.

Liu, Y., Murray, S.O., and Jagadeesh, B. (2009). Time course and stimulus dependence of repetition-induced response suppression in inferotemporal cortex. J Neurophysiol 101, 418–436.

Lorenc, E.S., Sreenivasan, K.K., Nee, D.E., Vandenbroucke, A.R.E., and D’Esposito, M. (2018). Flexible Coding of Visual Working Memory Representations during Distraction. J Neurosci 38, 5267–5276.

Lundqvist, M., Herman, P., and Miller, E.K. (2018a). Working Memory: Delay Activity, Yes! Persistent Activity? Maybe Not. J Neurosci 38, 7013–7019.

Lundqvist, M., Herman, P., Warden, M.R., Brincat, S.L., and Miller, E.K. (2018b). Gamma and beta bursts during working memory readout suggest roles in its volitional control. Nat Commun 9, 394.

Lundqvist, M., Rose, J., Herman, P., Brincat, S.L., Buschman, T.J., and Miller, E.K. (2016). Gamma and Beta Bursts Underlie Working Memory. Neuron 90, 152–164.

Meyer, T., and Olson, C.R. (2011). Statistical learning of visual transitions in monkey inferotemporal cortex. Proc Natl Acad Sci U S A 108, 19401–19406.

Mongillo, G., Barak, O., and Tsodyks, M. (2008). Synaptic theory of working memory. Science (New York, NY) 319, 1543–1546.

Muller, J.R., Metha, A.B., Krauskopf, J., and Lennie, P. (1999). Rapid adaptation in visual cortex to the structure of images. Science 285, 1405–1408.

Nikolić, D., Hausler, S., Singer, W., and Maass, W. (2009). Distributed fading memory for stimulus properties in the primary visual cortex. PLoS Biol 7, e1000260.

Parras, G.G., Nieto-Diego, J., Carbajal, G.V., Valdes-Baizabal, C., Escera, C., and Malmierca, M.S. (2017). Neurons along the auditory pathway exhibit a hierarchical organization of prediction error. Nat Commun 8, 2148.

Pasternak, T., and Greenlee, M.W. (2005). Working memory in primate sensory systems. Nat Rev Neurosci 6, 97–107.

Patterson, C.A., Wissig, S.C., and Kohn, A. (2013). Distinct effects of brief and prolonged adaptation on orientation tuning in primary visual cortex. J Neurosci 33, 532–543.

Peter, A., Uran, C., Klon-Lipok, J., Roese, R., van Stijn, S., Barnes, W., Dowdall, J.R., Singer, W., Fries, P., and Vinck, M. (2019). Surface color and predictability determine contextual modulation of V1 firing and gamma oscillations. Elife 8.

Priebe, N.J., Churchland, M.M., and Lisberger, S.G. (2002). Constraints on the source of short-term motion adaptation in macaque area MT. I. the role of input and intrinsic mechanisms. J Neurophysiol 88, 354–369.

Rademaker, R.L., Chunharas, C., and Serences, J.T. (2019). Coexisting representations of sensory and mnemonic information in human visual cortex. Nature neuroscience 22, 1336–1344.

Rademaker, R.L., van de Ven, V.G., Tong, F., and Sack, A.T. (2017). The impact of early visual cortex transcranial magnetic stimulation on visual working memory precision and guess rate. PLOS ONE 12, e0175230.

Rao, R.P., and Ballard, D.H. (1999). Predictive coding in the visual cortex: a functional interpretation of some extra-classical receptive-field effects. Nature neuroscience 2, 79–87.

Reinhart, R.M.G., and Nguyen, J.A. (2019). Working memory revived in older adults by synchronizing rhythmic brain circuits. Nat Neurosci 22, 820–827.

Romo, R., Brody, C.D., Hernandez, A., and Lemus, L. (1999). Neuronal correlates of parametric working memory in the prefrontal cortex. Nature 399, 470–473.

Rose, N.S., LaRocque, J.J., Riggall, A.C., Gosseries, O., Starrett, M.J., Meyering, E.E., and Postle, B.R. (2016). Reactivation of latent working memories with transcranial magnetic stimulation. Science (New York, NY) 354, 1136–1139.

Schwiedrzik, C.M., and Freiwald, W.A. (2017). High-Level Prediction Signals in a Low-Level Area of the Macaque Face-Processing Hierarchy. Neuron 96, 89–97 e84.

Scimeca, J.M., Kiyonaga, A., and D’Esposito, M. (2018). Reaffirming the Sensory Recruitment Account of Working Memory. Trends in cognitive sciences 22, 190–192.

Serences, J.T. (2016). Neural mechanisms of information storage in visual short-term memory. Vision research 128, 53–67.

Serences, J.T., Ester, E.F., Vogel, E.K., and Awh, E. (2009). Stimulus-specific delay activity in human primary visual cortex. Psychol Sci 20, 207–214.

Singer, W. (2021). Recurrent dynamics in the cerebral cortex: Integration of sensory evidence with stored knowledge. Proc Natl Acad Sci U S A 118.

Singer, W., and Lazar, A. (2016). Does the cerebral cortex exploit high-dimensional, non-linear dynamics for information processing? Frontiers in computational neuroscience 10, 99.

Sreenivasan, K.K., Curtis, C.E., and D’Esposito, M. (2014). Revisiting the role of persistent neural activity during working memory. Trends Cogn Sci 18, 82–89.

Stokes, M.G. (2015). ’Activity-silent’ working memory in prefrontal cortex: a dynamic coding framework. Trends Cogn Sci 19, 394–405.

Sugase-Miyamoto, Y., Liu, Z., Wiener, M.C., Optican, L.M., and Richmond, B.J. (2008). Short-term memory trace in rapidly adapting synapses of inferior temporal cortex. PLoS Comput Biol 4, e1000073.

Summerfield, C., and de Lange, F.P. (2014). Expectation in perceptual decision making: neural and computational mechanisms. Nat Rev Neurosci 15, 745–756.

Supèr, H., Spekreijse, H., and Lamme, V.A. (2001). A neural correlate of working memory in the monkey primary visual cortex. Science 293, 120–124.

Todorovic, A., and de Lange, F.P. (2012). Repetition suppression and expectation suppression are dissociable in time in early auditory evoked fields. J Neurosci 32, 13389–13395.

Trubutschek, D., Marti, S., Ueberschar, H., and Dehaene, S. (2019). Probing the limits of activity-silent non-conscious working memory. Proc Natl Acad Sci U S A 116, 14358–14367.

Uran, C., Peter, A., Lazar, A., Barnes, W., Klon-Lipok, J., Shapcott, K.A., Roese, R., Fries, P., Singer, W., and Vinck, M. (2022). Predictive coding of natural images by V1 firing rates and rhythmic synchronization. Neuron 110, 1240–1257 e1248.

van Kerkoerle, T., Self, M.W., and Roelfsema, P.R. (2017). Layer-specificity in the effects of attention and working memory on activity in primary visual cortex. Nat Commun 8, 13804.

Vinck, M., and Bosman, C.A. (2016). More Gamma More Predictions: Gamma-Synchronization as a Key Mechanism for Efficient Integration of Classical Receptive Field Inputs with Surround Predictions. Front Syst Neurosci 10, 35.

Vinje, W.E., and Gallant, J.L. (2000). Sparse coding and decorrelation in primary visual cortex during natural vision. Science 287, 1273–1276.

Wacongne, C., Labyt, E., van Wassenhove, V., Bekinschtein, T., Naccache, L., and Dehaene, S. (2011). Evidence for a hierarchy of predictions and prediction errors in human cortex. Proc Natl Acad Sci U S A 108, 20754–20759.

Wolff, M.J., Ding, J., Myers, N.E., and Stokes, M.G. (2015). Revealing hidden states in visual working memory using electroencephalography. Frontiers in Systems Neuroscience 9, 123.

Wolff, M.J., Jochim, J., Akyürek, E.G., and Stokes, M.G. (2017). Dynamic hidden states underlying working-memory-guided behavior. Nature Neuroscience 20, 864–871.

Yiling, Y., Shapcott, K., Peter, A., Klon-Lipok, J., Xuhui, H., Lazar, A., and Singer, W. (2023). Robust encoding of natural stimuli by neuronal response sequences in monkey visual cortex. Nat Commun 14, 3021.

